# Evaluation of methods to detect shifts in directional selection at the genome scale

**DOI:** 10.1101/2022.06.22.497174

**Authors:** Louis Duchemin, Vincent Lanore, Philippe Veber, Bastien Boussau

## Abstract

Identifying the footprints of selection in coding sequences can inform about the importance and function of individual sites. Analyses of the ratio of non-synonymous to synonymous substitutions (*d*_*N*_ */d*_*S*_) have been widely used to pinpoint changes in the intensity of selection, but cannot distinguish them from changes in the direction of selection, *i*.*e*., changes in the fitness of specific amino acids at a given position. A few methods that rely on amino acid profiles to detect changes in directional selection have been designed, but their performance have not been well characterized. In this paper, we investigate the performance of 6 of these methods. We evaluate them on simulations along empirical phylogenies in which transition events have been annotated, and compare their ability to detect sites that have undergone changes in the direction or intensity of selection to that of a widely used *d*_*N*_ */d*_*S*_ approach, `codeml`’s branch-site model A. We show that all methods have reduced performance in the presence of biased gene conversion but not CpG hypermutability. The best profile method, Pelican, a new implementation of [Tamuri et al., 2009], performs as well as `codeml` in a range of conditions except for detecting relaxations of selection, and performs better when tree length increases, or in the presence of persistent positive selection. It is fast, enabling genome-scale searches for site-wise changes in the direction of selection associated with phenotypic changes.

## 1 Introduction

The genomes and phenotypes of extant species bear traces of past adaptations that occurred in their ancestors. A lot of research in molecular evolution has been devoted to detecting and interpreting these traces, both in non-coding and coding sequences (e.g., [Moretti et al., 2014, Zhang et al., 2014, Merényi et al., 2020, Partha et al., 2019, Marcovitz et al., 2019]). In protein-coding genes in particular, several approaches have been developed to study evolution at the level of whole genes or at the level of single sites [Goldman and Yang, 1994, Yang and Nielsen, 2008]. Studies have found that amino acid changes at a single position could create an active site *de novo* [Risso et al., 2017], that amino acid changes at a few positions could change the affinity of an hormone receptor for its ligand [Bridgham et al., 2006], that convergent evolution could be detected at single sites in proteins in mammals [Li et al., 2010], in grasses [Christin et al., 2007], in insects [Zhen et al., 2012], and that amino acid changes at a single position could alter the dynamic of a worldwide viral epidemic [Korber et al., 2020]. Identifying traces of past and current adaptations at the level of single amino acid sites can thus be very insightful. In this article, we investigate the performance of several methods aiming to do just that. These include one commonly-used *d*_*N*_ */d*_*S*_ method, but also methods that have been more recently developed, based on *amino acid fitness profiles*.

In proteins, amino acids that are never or seldom encountered at a particular site in a group of related species may have been selected against in the past. Those that are frequent may have been favored by selection. One can study these differences in frequency to infer differences in fitness between amino acids. A *fitness profile* is then used to represent the relative fitness of each amino acid at a given site (fig. 1a: A, B, C and C’). When used within models of sequence evolution, a fitness profile determines the fixation probability of arising mutations during the process of evolution through mutation and selection [Halpern and Bruno, 1998, Yang and Nielsen, 2008, Rodrigue et al., 2010, Tamuri et al., 2012]. It also provides a *direction* for selection, which pushes evolution at the site away from low-fitness amino acids and towards high-fitness amino acids.

**Figure 1:**
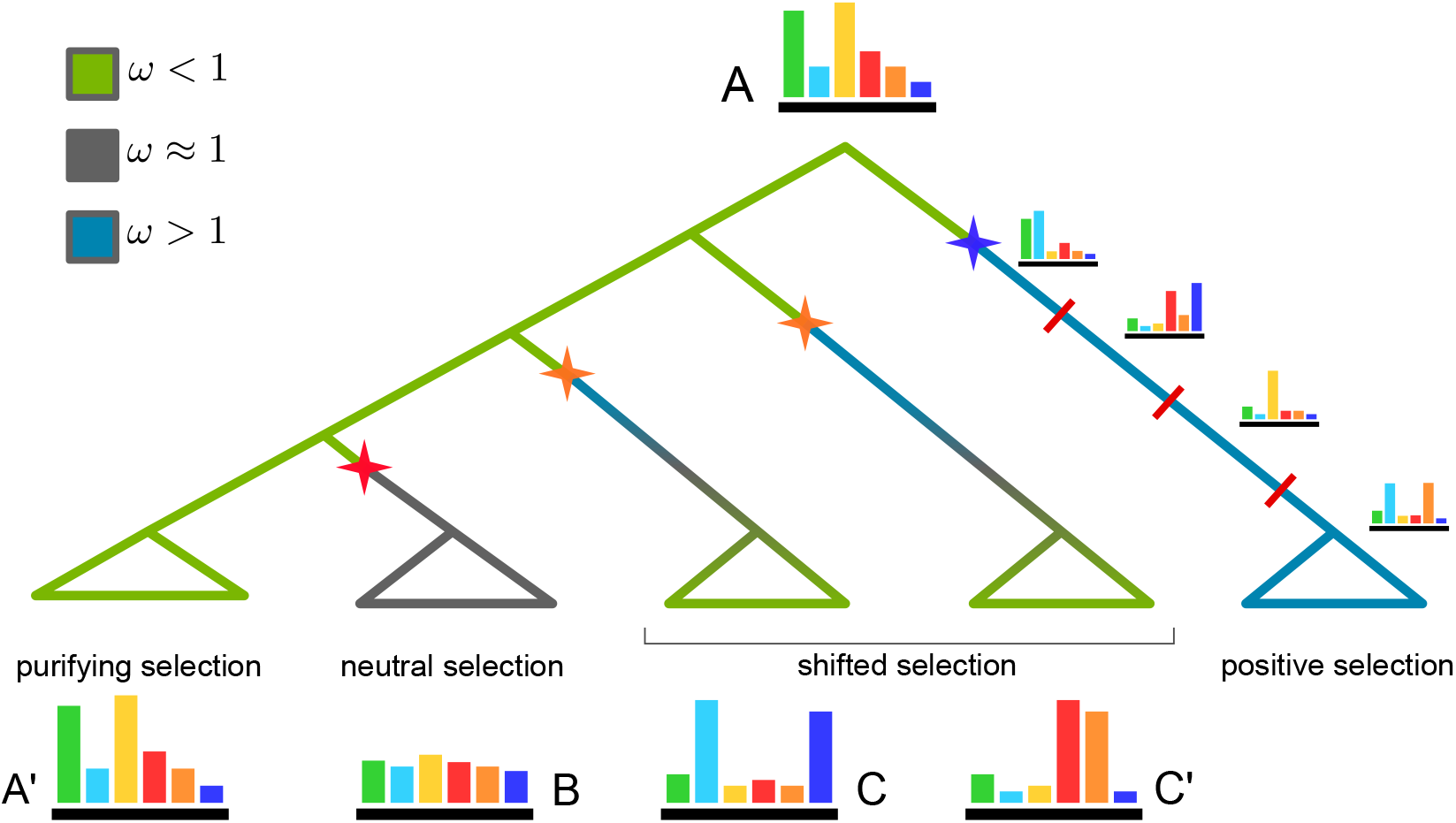
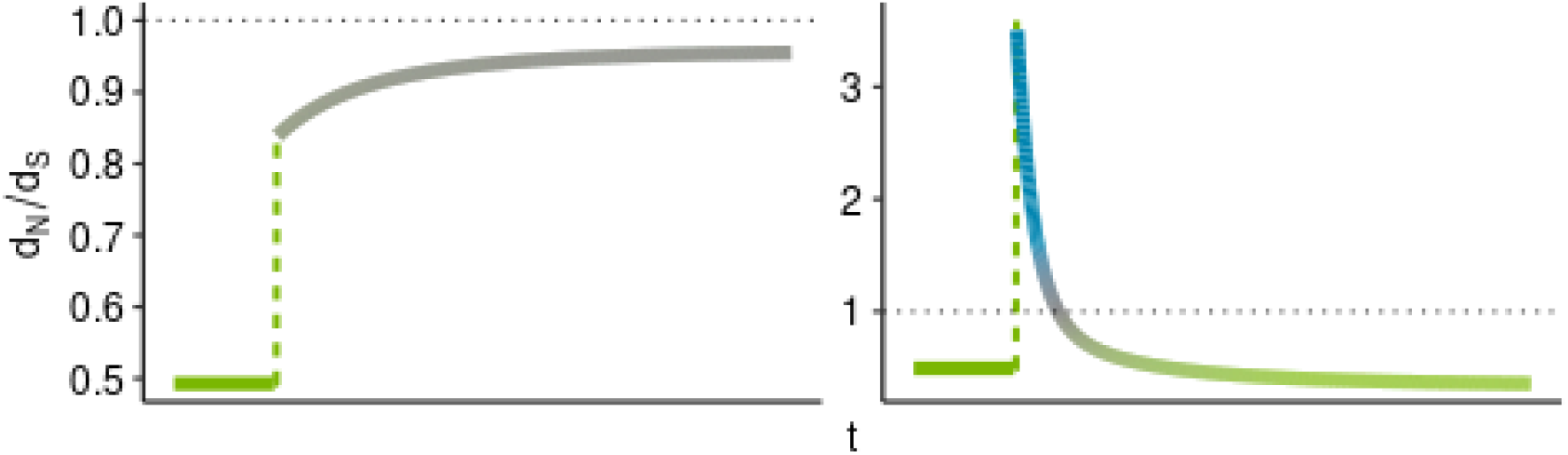
Schematic representation of various evolution scenarii of a protein site involving profile changes. Colored stars indicate transition events that trigger profile changes. The color gradient along branches show the variation of *d*_*N*_ */d*_*S*_ a.k.a *ω* values. The green sub-tree is a case of purifying selection, with fixed profile (A) and *ω <* 1. The grey sub-tree illustrates relaxed pressure subsequent to the transition in red, resulting in a flattened profile B and *ω ≈* 1. Two cases of shifted selection are represented, each one driven by a different fitness profile (C and C’). In both cases, there is a transient increase in the value of *ω*, followed by a decrease towards *ω <* 1, as represented on the right panel in 1b. Blue sub-tree is an example of persistent positive selection [Tamuri and dos Reis, 2021], where the fitness profile rapidly changes along the branch, at intervals marked with red bars. In this case, the value of *d*_*N*_ */d*_*S*_ remains greater than 1 while positive selection continues. (a) Cases of selection regime changes, with their representation as fitness Schematic representation of various evolution profile changes, and their equivalence with the *d*_*N*_ */d*_*S*_ metric. (b) *d*_*N*_ */d*_*S*_ variations over time. The curve on the left represents the simulated value of *d*_*N*_ */d*_*S*_ when transition from purifying selection to relaxed selection occurs (transition between profiles A and B above). On the right is the variation of *d*_*N*_ */d*_*S*_ during a shift in selection direction (transition between profiles A and C or C’ above).

The shape of a fitness profile derives from selective pressures that operate at a particular site of a protein. These pressures can be related to phenotypic traits or environmental constraints, which could change over time. In such a case, the pressures would change, and so would the fitness profile. Selection may vary in intensity, for instance as a trait becomes more or less important for the global fitness of the organism; and in direction, when changing the value of a trait leads to higher fitness. These different kinds of changes in selective pressure can be captured by variations of the fitness profile: changes in intensity through the pointedness or flatness of the profile (fig. 1a, transition from profile A to profile B), and changes in direction through the variation of the overall shape of the profile (fig. 1a, transition from profile A to profiles C and C’). In this manuscript we will focus on trait changes and the associated fitness profile at a site that occur discretely, at once, but progressive, continuous changes certainly occur in nature and would be important to consider.

Approaches to detect variations of selection on single sites of protein-coding sequences all require an annotation of the branches of a phylogeny, whereby each branch is associated to a phenotypic state or environmental condition. Given this annotation, either *d*_*N*_ */d*_*S*_ or profile methods can be used (fig. 2).

**Figure 2:**
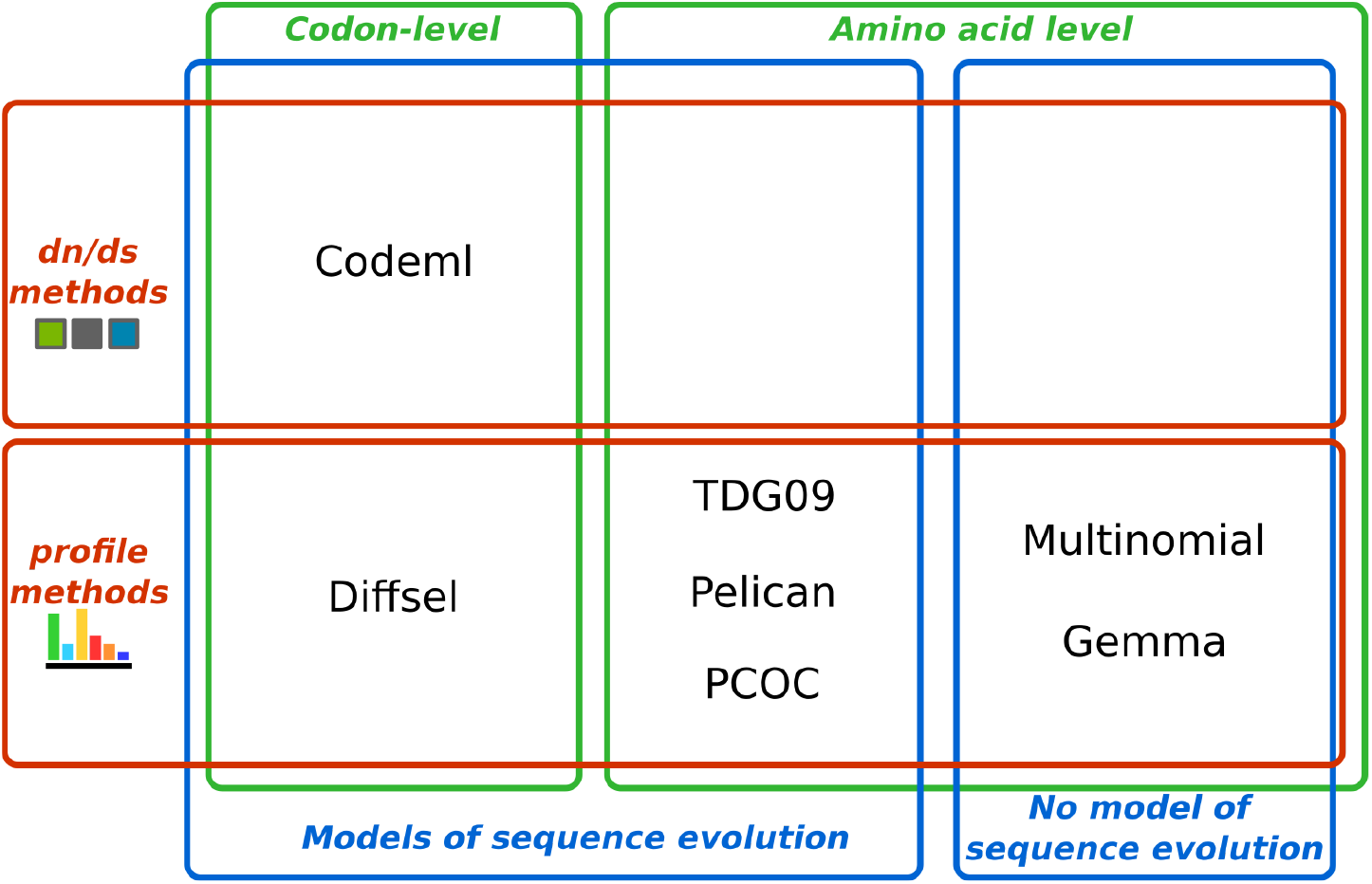
Methods evaluated in the manuscript. Methods have been positioned based on whether they are based on *d*_*N*_ */d*_*S*_ or amino acid profiles, whether they work at the codon or amino acid level, and whether they rely on a model of sequence evolution running along a phylogeny or not.

Approaches relying on the *ω* = *d*_*N*_ */d*_*S*_ metric have been widely used to capture variations in selective pressure [Kosiol and Anisimova, 2019], including in the context of genome screening (e.g. [Nielsen et al., 2005, Kosiol et al., 2008, Studer et al., 2008, Moretti et al., 2014, Zhang et al., 2014]). These methods can show good reliability, either at the level of whole gene sequences or of single sites, when the generating process matches the inference model (e.g., [Zhang, 2005]). The *ω* metric is defined as the ratio of rates between non-synonymous (*d*_*N*_) and synonymous (*d*_*S*_) substitutions. The underlying assumption is that selection operates at the amino acid level, so that synonymous codons provide the same fitness, while non-synonymous substitutions induce a variation in fitness. Inference is typically performed at the level of a gene by comparing the likelihood of a model with one set of *d*_*N*_ */d*_*S*_ values per condition, against a model having one global set of *d*_*N*_ */d*_*S*_ values through a likelihood ratio test (LRT) [Yang, 2007]. At the site level, the gene-wise parameter estimates are used to identify sites whose *d*_*N*_ */d*_*S*_ has changed in a manner correlated with the annotation of the phylogeny [Yang, 2005]. However, many other implementations have been proposed (e.g., [Guindon et al., 2004, Kosakovsky Pond et al., 2011, Murrell et al., 2015]). *d*_*N*_ */d*_*S*_ methods should be particularly effective at inferring changes in the intensity of negative selection: weaker (respectively stronger) selection should result in higher (resp. lower) *d*_*N*_ */d*_*S*_ values. In that sense, *d*_*N*_ */d*_*S*_ values have been used as a proxy of selection efficiency, even though in some cases this can be misleading [Spielman and Wilke, 2015, Jones et al., 2019]. In addition, *d*_*N*_ */d*_*S*_ methods should have good power to detect cases of persistent positive selection (right-most branch, fig. 1a), which should result in high *d*_*N*_ */d*_*S*_ values. However, they might be less effective at detecting changes in the direction of selection [Parto and Lartillot, 2018], as they may fail to detect some sites that have undergone episodic changes in directional selection on top of a background of strong purifying selection (see fig. 1 and [dos Reis, 2015]). Further, they do not output estimates of the direction of selection, but only *d*_*N*_ */d*_*S*_ values.

Profile methods have been developed more recently than *d*_*N*_ */d*_*S*_ methods, and have yet to be used at a genomic scale. They all rely on amino acid profiles to identify sites that correlate with a phenotype along a phylogeny, but vary in the complexity of their underlying models. Some methods operate at the codon level and can explicitly use amino acid fitness profiles by distinguishing between the mutation process, operating at the nucleotide level, and the selection process operating at the amino acid level (e.g., [Murrell et al., 2012, Parto and Lartillot, 2018]). These methods build on the *mutsel* framework [Halpern and Bruno, 1998, Yang and Nielsen, 2008, Rodrigue et al., 2010, Tamuri et al., 2012, Bloom, 2014] that provides a better description of coding sequence evolution than *d*_*N*_ */d*_*S*_ approaches [Spielman and Wilke, 2016, Bloom, 2014]. Other methods operate at the amino acid level and thus cannot model the mutation process. They use amino acid *frequency* profiles as a proxy to *fitness* profiles [Tamuri et al., 2009], and may thus be less powerful than methods that operate at the codon level. In both cases, inference can be performed with a Likelihood Ratio Test (LRT) at the site level, comparing the likelihood of a model with one profile per condition, against a model having one single profile that applies on all branches of the phylogeny. However, Bayesian inference has also been used, notably for a branch-heterogeneous mutsel model [Parto and Lartillot, 2018].

Both *d*_*N*_ */d*_*S*_ and profile methods to detect changes in selective pressures could be misled by non-adaptive processes, or by confounding between different selection regimes [Jones et al., 2019]. Non-adaptive processes notably include GC-biased gene conversion (gBGC) [Ratnakumar et al., 2010] [Bolívar et al., 2019] and CpG hypermutability [Meunier et al., 2005]. gBGC occurs during recombination and mimics natural selection by favoring the fixation of G and C alleles. CpG hypermutability increases the mutation rate of CG dinucleotides. These processes can generate patterns in the sequence data that could lead to false positives or false negatives, as has been shown for *d*_*N*_ */d*_*S*_ methods with respect to both gBGC [Ratnakumar et al., 2010, Guéguen and Duret, 2018, Rousselle et al., 2019], and CpG hypermutability [Saunders and Green, 2007, Suzuki et al., 2009]. Confounding between different selection regimes could happen if a test aiming to find changes in the direction of selection detected sites under persistent positive selection. It is unclear how sensitive profile methods would be to these problems.

Genome-scale detection of changes in selective pressure requires a fast method, especially if a large number of species is used to increase the power of an analysis. In fact, it has been suggested that the high computational cost of *d*_*N*_ */d*_*S*_ methods may be a hurdle to their more widespread use [Davydov et al., 2019]. It is unclear how efficient profile methods could be.

In this article, we evaluate several profile and *d*_*N*_ */d*_*S*_ methods to detect changes in selective pressures operating on individual positions of a protein-coding gene, on specific branches of a phylogeny. We consider several profile methods that have been published or that we have developed *de novo*, and compare them to a widely-used *d*_*N*_ */d*_*S*_ method. In particular, we ask whether profile methods can be as powerful as the *d*_*N*_ */d*_*S*_ method, including in the presence confounding factors, and pay particular attention to the computational costs of all methods.

Performance measurements are done on simulated datasets, allowing us to characterize the behaviour of the methods on a range of tree shapes, branch lengths, and number of transitions along the phylogeny. We also investigate whether the detection methods are sensitive to confounding signal generated by non-adaptive processes of molecular evolution [Ratnakumar et al., 2010, Bolívar et al., 2019, Meunier et al., 2005], or by persistent positive selection [Tamuri and dos Reis, 2021].

## 2 New Approaches

In this article, we introduce Pelican, an improved implementation of the model from [Tamuri et al., 2009]. This implementation was found to have better sensitivity and specificity than the original, and is also faster thanks to optimisations on linear algebra computation.

Multinomial is a fast non-phylogenetic profile method that is also evaluated in this paper. It models observed amino acid frequency profiles as multinomial distributions, and compares the likelihoods at a given site of a single frequency profile versus multiple profiles through a likelihood ratio test.

Both of these methods are implemented as a single program, that is made available to detect differential selection in protein sequence alignments. In this context, Multinomial can be used as a fast filter on the alignment to reduce the amount of candidate sites to be evaluated through Pelican.

## 3 Results

We evaluated the performance of detection methods using simulated datasets. The methods that were considered are represented in fig. 2 and include :

- `codeml`, a widely used *d*_*N*_ */d*_*S*_ method for detecting positive selection, provided in the PAML toolkit [Yang, 2007]. `codeml` was configured to use the branch-site model A [Zhang, 2005, Yang, 2005], and works at the codon level.
- Multinomial, the simplest and fastest profile method, does not rely on a model of sequence evolution and works at the amino acid level. It uses a likelihood ratio test to compare two models, one in which a single amino acid profile is used to describe amino acid frequencies observed at a site across all tip sequences, and one where different amino acid profiles are used depending on the condition associated to the tip. Multinomial ignores the shape of the phylogeny and could thus be misled by phylogenetic inertia.
- Gemma [Zhou and Stephens, 2012], based on a linear mixed model, was originally developed for genome-wide association studies (GWAS). It does not use a model of sequence evolution, but can take into account the structure of the phylogeny, encoded as a correlation matrix, which is introduced as a random effect in the mixed model. We used it at the amino acid level, by encoding the protein alignment as an alignment of binary characters (see section 6).
- TDG09 [Tamuri et al., 2009], a profile method that can be considered as a refinement over the Multinomial method, in that it also works at the amino acid level but takes into account the phylogeny by relying on a model of sequence evolution. It uses a LRT to compare a model with one profile per condition and a model with one single global profile.
- Pelican, a new implementation of the model underlying TDG09 [Tamuri et al., 2009], originally motivated by the observed discrepancy reported between the performances of Diffsel and TDG09.
- PCOC [Rey et al., 2018], a profile method working at the amino acid level. It is at its base similar to TDG09 but works with a limited set of pre-existing profiles, and further expects to observe substitutions at every transition between conditions in the phylogeny.
- Diffsel [Parto and Lartillot, 2018], a profile method working at the codon level and based on a mutation-selection model in a Bayesian framework. Diffsel has performed significantly better than the other methods in a previous benchmark [Rey et al., 2019].

All simulations were done under a codon-based, time-reversible, mutation-selection model with site-specific amino acid fitness profiles. The model was run along a phylogeny whose branches are annotated with two conditions that we refer to as *background* and *foreground*. A simulation generates codon and corresponding amino acid alignments of arbitrary length. Sites in the alignment may be either: (1) *H*_*A*_ sites, that are the result of a simulation where changes in the selection dynamic occur between background and foreground branches ; (2) *H*_0_ sites, resulting from an evolutionary process where selection is constant. The number of sites of each type was controlled in the simulation, allowing the comparison of predictions on the nature of each site (*H*_0_ or *H*_*A*_) with its known type, to estimate the performances of the prediction method.

Performance estimates in all the benchmarks were done using two metrics : *precision* and *recall*. Precision is the proportion of true positive sites among all sites identified as positive. Recall, also known as sensitivity, is the proportion of *H*_*A*_ sites that are identified as positive. These metrics were summarized by computing the area under the precision-recall curve (PR AUC). Confidence intervals for the PR AUC were computed according to [Boyd et al., 2013].

In this section we compared the detection methods using our simulation model in several contexts : (1) synthetic trees with variable branch lengths and numbers of transitions; (2) empirical trees in the presence or absence of confounding factors in the simulation. In the following, all branch length values are given in expected numbers of codon substitutions.

### 3.1 Detection performances on synthetic trees

#### 3.1.1 Detection performances increase with the number of transitions

We investigated whether the number of transitions from background to foreground conditions had an effect on detection performances. We generated a balanced tree of 128 tips in which all branch lengths equal 0.01, and generated a variable number of transitions on terminal branches (tree topology shown in sup. fig. S1). In this setting, both the number of foreground leaves and the total time in the foreground condition increase with the number of transitions. Results shown in fig. 3a show that all methods take advantage from such increases.

**Figure 3:**
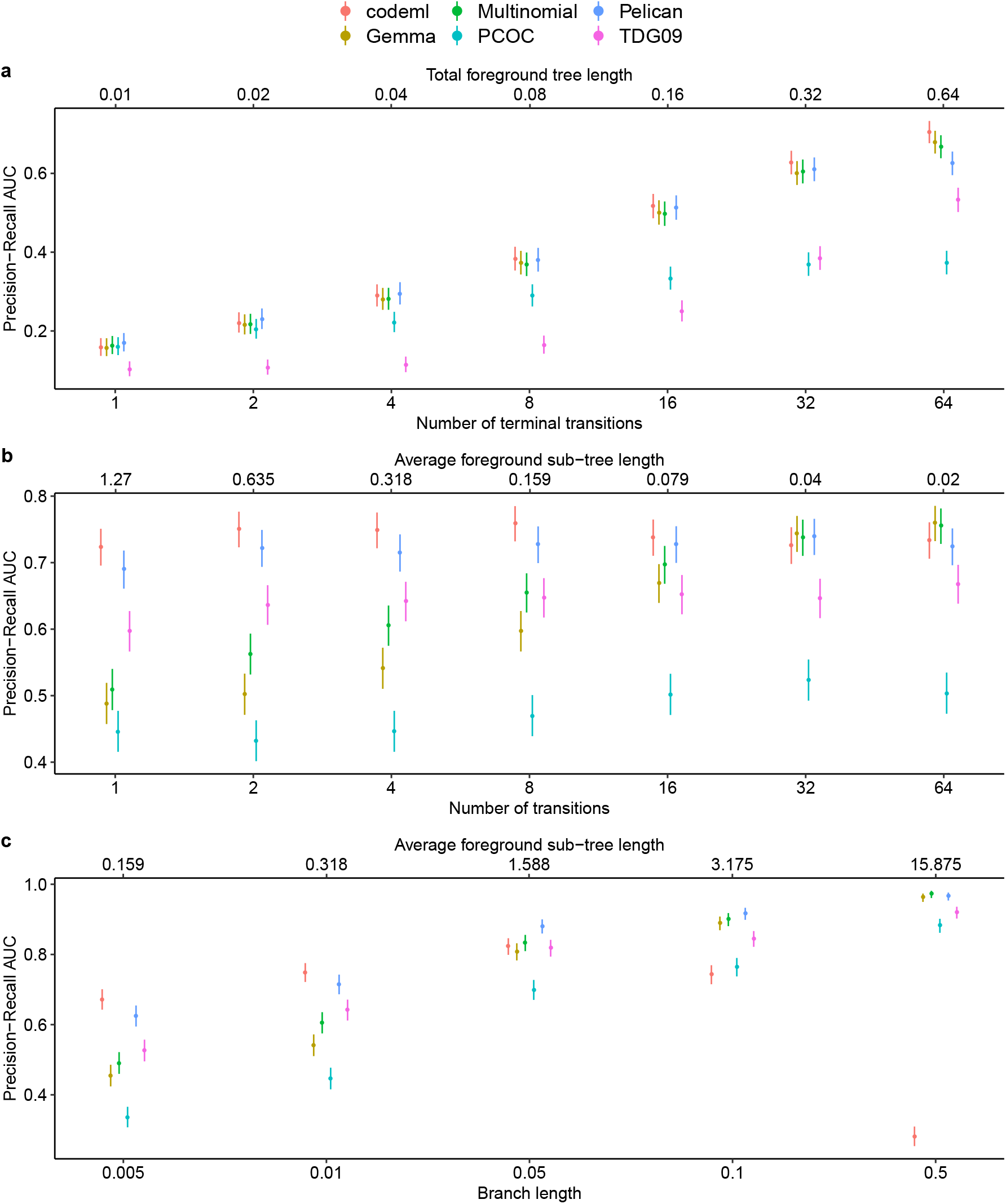
Detection performances evaluated on synthetic trees. 95% confidence intervals accounting for the variability of the PR AUC estimates are shown. (a) Performance increases with the number of transitions on terminal branches. (b) The number of transitions is not the determining factor for the performance of the phylogenetic methods but has a strong effect on the performance of Gemma and Multinomial. (c) Performances of the profile methods are positively correlated to the branch lengths, while the performance of `codeml` decreases on longer branches.

#### 3.1.2 The amount of time spent in a condition has a large effect on detection performance for phylogenetic methods

We next evaluated the relative importance of the number of transitions and the amount of time spent in the foreground condition on the phylogeny. We used a different set of trees with the same general features (128 tips, branch lengths equal 0.01), varied the numbers of transitions, but kept the number of foreground leaves and total foreground length constant across trees. This was done by normalizing the branch lengths to achieve equal total times between foreground and background conditions, and across trees. As a result the number of tips in each foreground sub-tree is variable, depending on the depth of the transition event. For a given number of transitions, all transitions occur at the same depth in the tree (sup. fig. S2).

Figure 3b shows that the performances of Gemma and Multinomial increase with the number of transitions, even when the amount of time spent in the foreground condition is kept constant. They become the best performing methods at 64 transitions. However, the phylogenetic methods `codeml` and Pelican seem to be less sensitive to this parameter in this experiment, suggesting that the determining factors for their performances in the previous experiment were the total foreground time and/or the number of foreground leaves, which are kept constant in this experiment.

#### 3.1.3 Profile methods improve as branch lengths increase

In order to assess the effect of branch lengths and of the distance between transition events and foreground leaves on method accuracy, while keeping the number of transitions constant, we evaluated each method on a balanced tree with 4 transition events where a scaling factor was applied to the branch lengths (sup. fig. S3). As a side-effect, this scaling factor also applies to the total foreground tree length.

Results in figure 3c highlight two opposite trends between profile methods and the *d*_*N*_ */d*_*S*_ method `codeml`, in relation with the branch length scaling. Profile methods tend to be more accurate in detecting selection shifts when the branch lengths increase, while the performance of `codeml` decreases. We suspect that as branch lengths increase, the number of synonymous substitutions increases, which reduces *d*_*N*_ */d*_*S*_ and makes it harder to detect *H*_*A*_ sites (see fig. 1b, right).

Among profile methods, the performance gap tends to decrease with longer branches.

### 3.2 Detection performances on empirical phylogenies

To benchmark the methods in a more realistic context, we evaluated their performances on empirical phylogenies that differ in their size, depth and number of transitions (Table 1). The corresponding phylogenetic trees are shown as supplementary material (supplementary figures S4, S5, S6, S7, S8, S9).

**Table 1:** Summary statistics on empirical trees. Tree depth is defined here as the highest number of branches between a leaf and the root. Size is the number of leaves in the tree. Transitions are defined as changes from the background to the foreground condition.

| Dataset | Depth | Size | Transitions | Avg branch length |  | Avg sub-tree length<br>Foreground |
| --- | --- | --- | --- | --- | --- | --- |
|  |  |  |  | Global | Foreground |  |
| Rodents<br>[Rey et al., 2019] | 11 | 32 | 10 | 0.0192 | 0.0252 | 0.0353 |
| *Cyperaceae*<br>[Besnard et al., 2009] | 25 | 79 | 5 | 0.0207 | 0.0239 | 0.196 |
| Echolocation<br>[Scornavacca et al., 2019] | 18 | 116 | 3 | 0.0081 | 0.0061 | 0.0828 |
| *Amaranthaceae*<br>[Kapralov et al., 2012] | 22 | 179 | 15 | 0.0045 | 0.0035 | 0.0356 |
| HIV RTi<br>[Murrell et al., 2012] | 34 | 476 | 238 | 0.0063 | 0.0051 | 0.0051 |
| Influenza H1 segment<br>[Tamuri et al., 2009] | 61 | 434 | 1 | 0.0603 | 0.0608 | 49.0709 |

Alignments were simulated as in the previous experiments, using the simulation model running along the empirical phylogenies. These alignments were used to measure the statistical calibration and the throughput of each method, and to evaluate each method as in the previous section.

#### 3.2.1 Pelican performs well on empirical phylogenies

We assessed whether the methods were well calibrated, *i*.*e*., how accurate was their reported false positive rate under the null (H0) model. To this end, we simulated 9,000 sites under H0, and counted the number of false positives for each method, at the 0.05 p-value threshold. Under this setting, a well calibrated method should produce on average a number of false positives equal to 5% of the total number of sites. Results shown in sup. table S1 indicate that most methods are overly conservative, *i*.*e*., their observed false positive rate is lower than their advertised (5% here) false positive rate. Multinomial is the only method that can yield a higher rate of false positives, particularly on the Influenza phylogeny. To further assess how conservative the methods were, we computed the observed false positive rate on non-constant sites only, given that constant sites cannot be classified as positive. Sup. table S2 indicates that even on this subset of sites, most methods still have low rates of false negatives. This indicates that all methods except Multinomial are overly conservative.

We then assessed the performance of the methods to detect *H*_*A*_ sites by simulating 1000 *H*_*A*_ sites and 9000 H0 sites. Pelican, `codeml` and Diffsel consistently show the best performances on all datasets (fig. 4), with the exception of the Influenza H1 dataset. It is worth noting that Diffsel is one of the best performing methods, even though it estimates branch lengths and does not get them as input, like most other methods.

**Figure 4:**
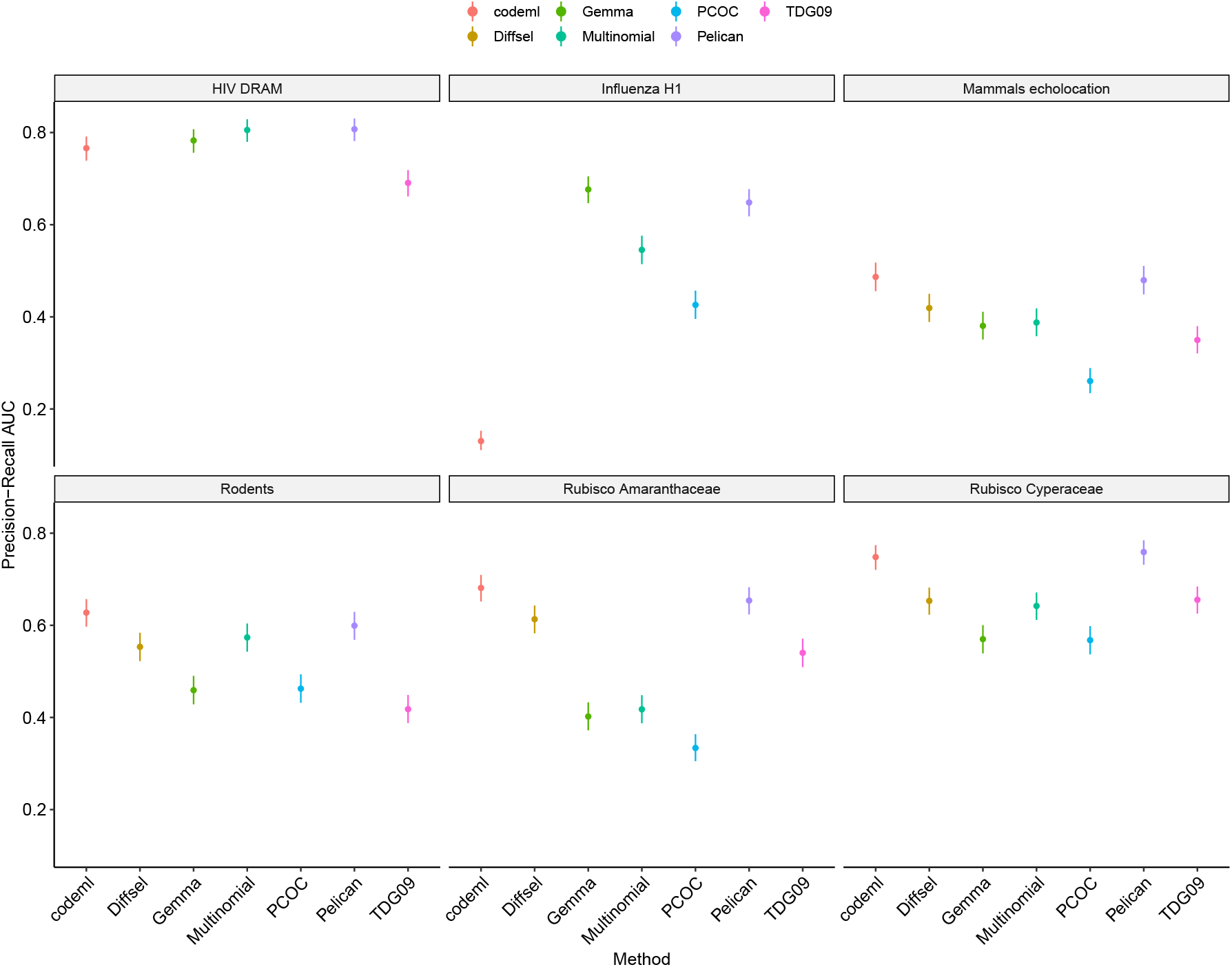
Precision-Recall area under the curve (AUC) estimates on simulated datasets using 6 empirical phylogenies, under changes in the direction of selection. Performances of TDG09 on the Influenza H1 dataset were not successfully measured. Diffsel was not evaluated on the HIV and Influenza dataset due to the large computation times involved. PCOC had an underflow error on the HIV data set.

We note that, while Pelican is essentially a reimplementation of TDG09, it shows significantly better performances on every dataset. `codeml` and Pelican have similar performances in general. However, on the Influenza H1 dataset, which has the highest average foreground sub-tree length 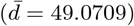, `codeml` incurs a large drop in its performances. These observations are consistent with the results obtained on synthetic trees (fig. 3c).

Even though the HIV dataset has the lowest average foreground sub-tree length (*d* = 0.0051), Pelican performs better than `codeml` on this dataset. Performances are strongly increased for all methods on this dataset, compared to the other empirical phylogenies. Our explanation for the results on the HIV dataset involves multiple effects : (1) the large number of transitions (*n* = 238) on terminal branches yields a strong signal for all methods, which benefits profile methods the most (see fig. 3a) ; (2) figure 3c seems to indicate that there is an optimal branch length for `codeml`: the signal for *d*_*N*_ */d*_*S*_ falls off on longer branches, but branches can also be too short to allow reliable *d*_*N*_ and *d*_*S*_ estimations because of the insufficient number of substitutions occurring in such a short time span.

We showed that some characteristics of the phylogenies had a major effect on method performance, particularly the time spent in the foreground condition, as well as the number of transitions in the phylogeny. It is likely that variations in the detection performances are the results of interactions between the features of the phylogeny, possibly including more than the two we identified, as well as the sensitivity of the detection method to these features.

On a side note, we remark that Multinomial shows some surprisingly good performances despite its simplicity. As it does not take any information from the phylogeny, it is the simplest profile method, and also the fastest (table 2).

**Table 2:** Execution times for one alignment containing 100 *H*_0_ and 100 *H*_*A*_ sites generated using our collection of empirical phylogenies. Result for TDG09 on the Influenza dataset is not available due to the program not terminating within a reasonable amount of time.

| Method | Execution time (s) |  |  |  |  |  |
| --- | --- | --- | --- | --- | --- | --- |
|  | *Cyperaceae* | *Amaranthaceae* | Rodents | Echolocation | HIV | Influenza |
| Multinomial | 0.01 | 0.02 | 0.01 | 0.02 | 0.04 | 0.03 |
| Gemma | 1.79 | 1.90 | 1.72 | 1.81 | 1.96 | 2.03 |
| Pelican | 10.93 | 19.42 | 2.58 | 6.72 | 87.72 | 266.84 |
| TDG09 | 22.21 | 40.56 | 6.84 | 12.56 | 369.87 |  |
| codeml | 60.60 | 172.50 | 27.37 | 100.76 | 443.66 | 614.33 |
| PCOC | 65.56 | 129.01 | 27.80 | 76.48 | 346.25 | 436.84 |
| DiffSel | 1253.00 | 1497.84 | 946.79 | 1083.48 | 2659.78 | 3982.00 |

#### 3.2.2 Performances in the detection of changes in the intensity of selection

Profile methods are in principle particularly appropriate for detecting changes in the direction of selection, and in practice perform as well as `codeml` and better on long branches (see above).

We evaluated how they perform in the presence of a change in the intensity of selection, by simulating a scenario of relaxation of selection. In this scenario, *H*_*A*_ sites are simulated such that all amino acids have equal fitness on foreground branches. This corresponds to a complete relaxation of selection.

Fig. 5 indicates that profile methods, and Pelican in particular, can also detect relaxations of selection, but that their performance depends on the phylogeny. In particular, we find that in some cases the detection is unreliable (fig. 5, Influenza panel). We suspected that this lower performance was due to a lack of sensitivity, and tested this hypothesis by changing the computation of degrees of freedom in the Likelihood Ratio test performed in Pelican (sup. section S5). Sup. fig. S13 shows that much better performances can be obtained on the Influenza data set, but with some cost on the performance of the method on other data sets (notably Mammals echolocation). Future work on the LRT may result in an improved performance of Pelican across data sets, in settings of changes in both the direction and the intensity of selection.

**Figure 5:**
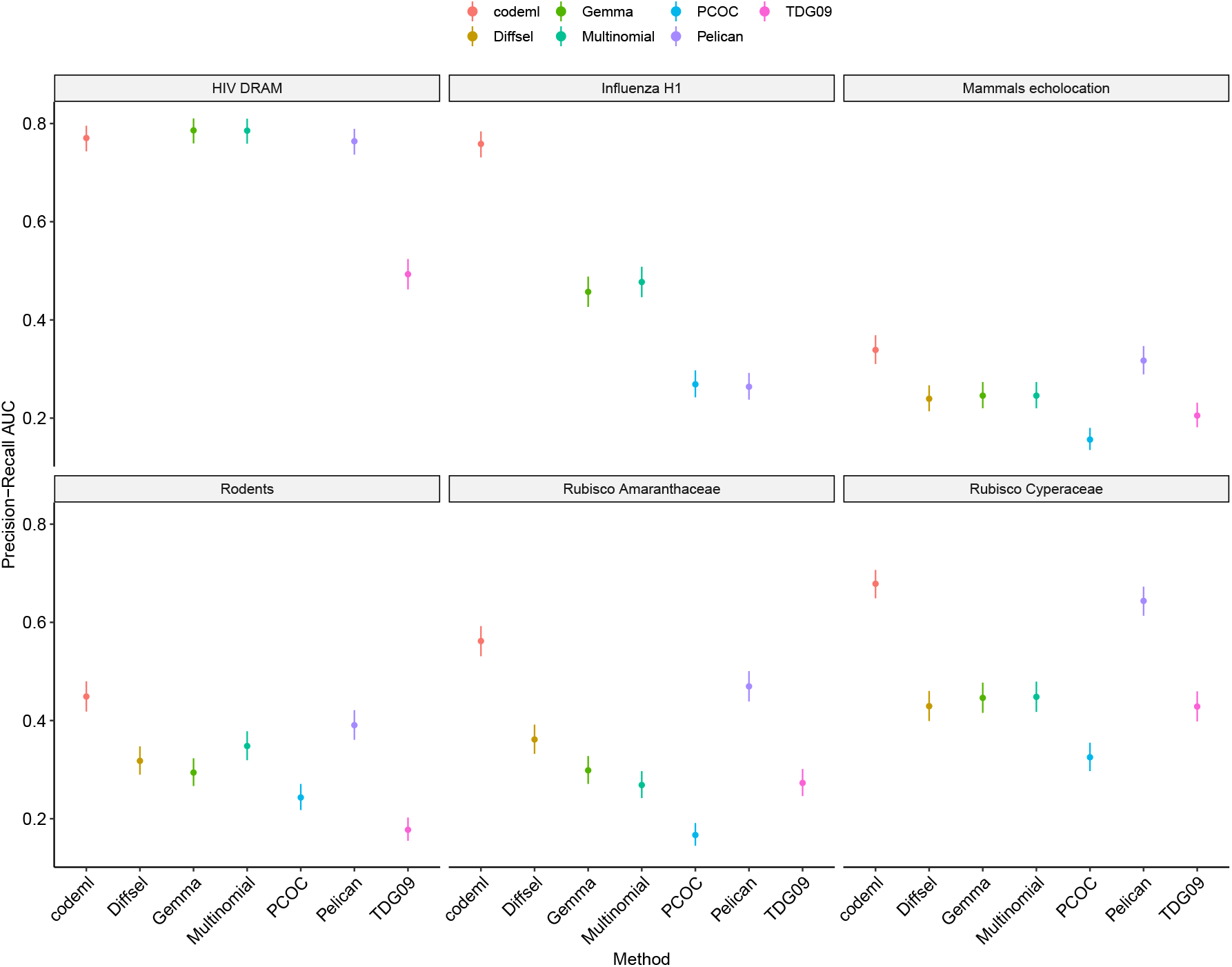
Precision-Recall area under the curve (AUC) estimates on simulated datasets using 6 empirical phylogenies, under relaxation of selection. Performances of TDG09 on the Influenza H1 dataset were not successfully measured. Diffsel was not evaluated on the HIV and Influenza dataset due to the large computation times involved. PCOC had an underflow error on the HIV data set.

#### 3.2.3 Performances in the presence of confounding factors

In order to assess the robustness of the detection to other evolutionary processes, we executed a benchmark on simulations including confounding factors: (1) CpG hypermutability, which induces a higher mutation rate on methylated CpG dinucleotides; (2) GC-biased gene conversion (gBGC), a non-adaptive process that increases the overall GC content in the genome and may be mistaken as a selective force [Ratnakumar et al., 2010]; (3) persistent positive selection (PPS), as modeled by [Tamuri and dos Reis, 2021], which favors non-synonymous substitutions over synonymous ones on the branches where it occurs. We used strong but realistic intensities for each of these processes, with two intensities for gBGC, and two intensities for PPS. In simulations of CpG hypermutability and GC-biased gene conversion (gBGC), the processes were applied on foreground branches for both H0 and HA sites. In the simulation of PPS, the process was applied on all branches, but only on H0 sites, to assess the propensity of each method to generate false positives. Results are shown in figure 6 for the Echolocation phylogeny, and are available as supplementary material for the other phylogenies.

**Figure 6:**
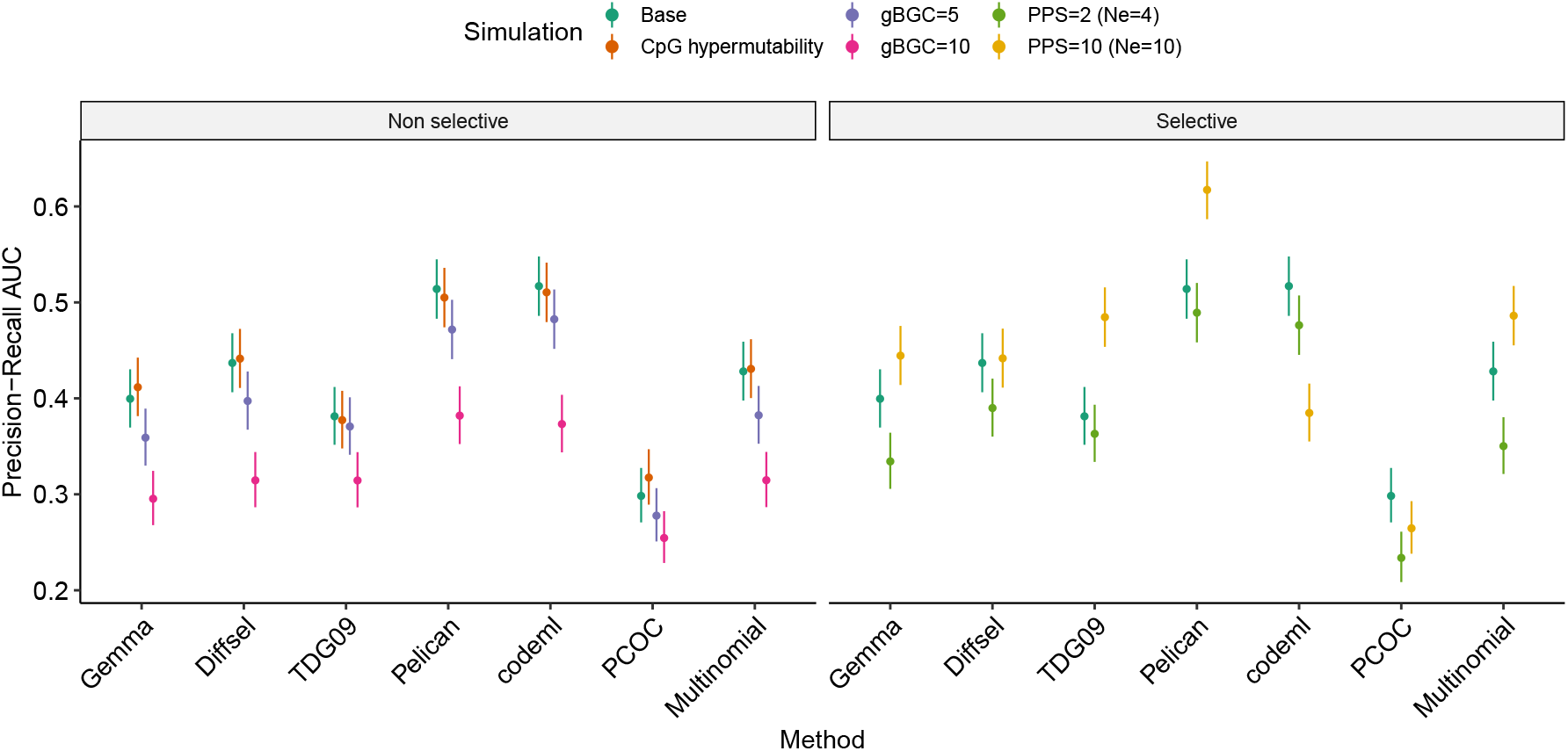
Effects of GC-biased gene conversion (gBGC), CpG hypermutability and persistent positive selection (PPS) on precision-recall AUC on the Echolocation dataset.

We find that the presence of CpG hypermutability has no influence on the detection performance in most cases.

In contrast, on simulations including gBGC, we notice a strong decrease of the performance for every method. While gBGC happens at the nucleotide level, it generates selection-like signal at the codon (or amino acid) level, that is not the result of an adaptive process. This signal was strong enough to directly interfere with the signal for selection on genomic sites.

At a fixed effective population size *N*_*e*_, an increase in PPS results in a decrease in the performance of all methods. Under conditions of strong PPS and large *N*_*e*_, the performance of `codeml` is strongly reduced, but the performance of profile methods can be improved. Overall, profile methods seem less prone to generating false positives in the presence of PPS, the effect of which is largely compensated by an increased value of *N*_*e*_.

#### 3.2.4 Throughput varies greatly between methods

We measured execution time for each method on six simulated datasets. Simulations were made using our simulation model on each empirical tree to generate an alignment of 100 *H*_0_ and 100 *H*_*A*_ sites. Execution times were measured as the elapsed time at completion of a run for each method using a single CPU, and are presented in table 2. The throughput of phylogenetic methods can vary by a large factor depending on the size of the phylogeny.

Multinomial and Gemma are the fastest methods by a large factor. None of these two methods require parameter estimations for a model of sequence evolution, allowing faster execution. At the other end, the two codon-level methods `codeml` and Diffsel are the slowest. Pelican is the fastest of the phylogenetic methods by a non negligible factor on all datasets.

## 4 Discussion

In this paper, we used simulations to compare the performance of methods that detect changes in the direction and intensity of selection, given an annotation of a phylogeny. These simulations rely on mutation-selection models of codon sequence evolution running along phylogenies.

### 4.1 Mutation-selection models for simulating coding sequences

Our choice to rely on mutation-selection models stems from the fact that these models have been shown to be more realistic for coding sequences than *d*_*N*_ */d*_*S*_ methods [Spielman and Wilke, 2016, Bloom, 2014]. They distinguish between processes occurring at the mutation level, and processes occurring at the selection level among codons. This flexibility allowed us to implement in our simulations CpG hypermutability and gBGC. In addition, we have made the choice to use site-heterogeneous amino acid fitness profiles to emulate the heterogeneity among positions in protein sequences. For improved realism, the profiles we used come from [Rey et al., 2019], and are based on laboratory mutagenesis experiments [Bloom, 2017]. However, we assumed no fitness difference between synonymous codons, even though this can be implemented in the mutation-selection framework [Yang and Nielsen, 2008, Pouyet et al., 2016]. Despite this, and given the fact that we simulated along empirical phylogenies, we expect our results are representative of the performance of the methods on empirical data sets.

### 4.2 Methods working at the amino acid level perform as well as codon-based methods

Some of the methods in the benchmark rely on models that are similar to our simulation model. In particular, Diffsel is also based on a mutation-selection model, and `codeml` works at the codon level. Expectedly, these two methods perform well on our simulations. In agreement with previous results [Spielman and Wilke, 2015], `codeml`, which relies on *d*_*N*_ */d*_*S*_ and does not use amino acid fitness profiles, is very effective except on long branches and trees (fig. 3c and fig. 4 the Influenza H1 phylogeny). All the other methods work at the amino acid level. Among those, the models based on a phylogenetic model (Pelican, TDG09, PCOC) vary in their performance, with Pelican standing out as the best performer. The lower performance of PCOC is likely due to two of its characteristics. Firstly, its reliance on a predefined set of amino acid frequency vectors, which may prevent it from accurately fitting the sites under study. Secondly, its “One-Change” component, which requires an amino acid change at each transition between background and foreground branches. This second limitation by design reduces the number of positive sites it can detect. TDG09 has lower performance than its reimplementation Pelican. The two implementations agree in the majority of cases, but disagree on some sites, likely due to optimization problems on boundary cases, which penalize TDG09. The fact that Pelican’s performance is similar to the performance of codon-based models suggests that its reliance on a WAG exchangeability matrix, not used in the simulation model, is not harmful. Further, it suggests that no information present only at the codon level is of much use to `codeml` or Diffsel, even when sequences are simulated with a model of CpG hypermutability (fig. 6). This may seem surprising, but probably relates to how we specified the detection problem we addressed. It is entirely centered around the amino acid profiles, so the codon level does not provide much useful information. Finally, the non-phylogenetic methods perform quite well despite their simplicity. Multinomial, the simplest of our methods, performs better than Gemma, which has the ability to include the shape of the phylogeny as a covariate. This may be because Gemma was designed to handle binary characters, and we had to transform the amino acid data before feeding it into Gemma (see methods).

Beyond detection efficacy, the *d*_*N*_ */d*_*S*_ and profile methods that we discuss in this manuscript vary in their execution speed. Methods that rely on models of sequence evolution typically have large computational footprints due to the use of the pruning or sum-product algorithm [Felsenstein, 1981], and the need for frequent matrix exponentiations. The computational footprints of these operations become larger as the state space grows: methods that work at the codon level (61 states) are more demanding than methods that work at the amino acid level (20 states) (fig. 2). Therefore, the profile methods that work at the amino acid level benefit from a computational advantage compared to codon-level profile methods or *d*_*N*_ */d*_*S*_ methods. Diffsel is the slowest method despite a thoroughly optimized code base, for several reasons. Firstly, it works at the codon level. Secondly, it attempts to estimate more parameters than the other methods, and notably branch lengths. Thirdly, it is the only Bayesian method here, and as such is the only one providing a credible interval for each parameter, at each position, where the other methods only provide point estimates. Pelican’s speed is better than TDG09’s, due to the reliance on high performance computing libraries (see methods). It also uses diagonalization for matrix exponentiations, or the contraction of sparse substitution matrices to matrices of lower sizes as in the original method [Tamuri et al., 2009]. It has thus already been extensively optimized, but further improvements might be obtained by using substitution mapping and summary statistics as in Diffsel [Parto and Lartillot, 2018].

### 4.3 Features of a data set that affect performances

Results obtained in this benchmark highlight that profile and *d*_*N*_ */d*_*S*_ methods perform differently in detecting changes in directional selection, depending on the features of a dataset. We identified a set of tree features that appear to have an effect on the performances: the number of transitions from background to foreground condition, the total time in the foreground condition, the number of foreground leaves, and the average length of foreground sub-trees. The variations in the detection performances observed on empirical phylogenies (fig. 4) likely are the result of interactions between these features, and possibly others that have yet to be identified.

Both Pelican and `codeml` benefit from an increased number of leaves in the foreground condition (fig. 3a), but not from an increased number of transitions (fig. 3b). However, non-phylogenetic methods (Multinomial, Gemma) benefit from increasing any of these features, including the number of transitions.

`codeml` tends to perform better than profile methods on phylogenies with shorter foreground sub-trees on average. This conforms to our understanding of the two types of methods, and the kind of signal they rely on, as presented in section 1. In the case of a change in the direction of selection, the resulting burst of the *d*_*N*_ */d*_*S*_ ratio occurs over a short time period, and quickly decreases back to a purifying selection regime (*d*_*N*_ */d*_*S*_ *<* 1, fig. 1b). This implies that on longer branches more time is spent in a purifying selection regime, reducing the signal for high *d*_*N*_ */d*_*S*_ as the rate of non-synonymous substitutions decreases.

In contrast, profile methods rely on amino-acid frequencies to detect positive selection. In this case, the signal is strongest when the amino-acid frequencies have reached an equilibrium and differ the most from the ancestral frequency distribution. As reaching the foreground equilibrium distribution through substitutions takes time, detection performances tend to increase on longer branches (fig. 3c).

Profile methods that do not take into account the phylogenetic information have a reduced performance on short branches. In that case observations at the leaves of the phylogenetic tree are more strongly correlated and this may mislead methods that assume independent observations (like Multinomial) or rely on a less accurate model (like Gemma). On longer branches, observations at the leaves of the tree tend to become more independent, and non phylogenetic methods exhibit performances similar to their more complex counterparts.

### 4.4 GC-biased gene conversion is an important confounding factor for both *d*_*N*_ */d*_*S*_ and profile methods

In an effort to make our simulations more realistic, we introduced two non-adaptive confounding factors in our model: CpG hypermutability, which affects the mutation component, and GC-biased conversion (gBGC), which affects the selection component. We have found that introducing gBGC on foreground branches induces a significant drop in performances for all the evaluated methods, with higher values of gBGC resulting in larger decreases (fig. 6). gBGC mimics selection, independently of the underlying fitness profiles, and scrambles the signal used to detect changes in the selection regime. This corroborates previous studies on the role of gBGC in disrupting the detection of selection in genome sequences [Ratnakumar et al., 2010, Rousselle et al., 2019, Guéguen and Duret, 2018]. Mechanistic codon-level models such as Diffsel could be extended to account for this effect, and untangle it from directional selection.

On the other hand, strong CpG hypermutability was not found to induce changes in the performance in most cases. It is possible that codons that contain CG dinucleotides are not frequent enough in our simulations based on the mutsel framework to reduce the AUC metric.

### 4.5 Persistent positive selection is an important confounding factor for *d*_*N*_ */d*_*S*_ methods, less so for profile methods

Protein sites may be subject to a variety of selection regimes (fig. 1a). It may be difficult to distinguish sites undergoing changes in the direction of selection from sites evolving under a different selection regime, in particular persistent positive selection. In our simulations under the model of [Tamuri and dos Reis, 2021], we found that `codeml` had difficulty distinguishing the two processes, in agreement with [Parto and Lartillot, 2018]. PPS results in elevated (*>* 1) *d*_*N*_ */d*_*S*_ values throughout the phylogeny, which is not well modelled by `codeml`’s branch-site model A, which assumes that positive selection only occurs on foreground branches. `codeml` has to choose between two alternatives, none of which fits the data very well: either consider that the PPS sites never have *d*_*N*_ */d*_*S*_ *>* 1, or consider that the PPS sites have *d*_*N*_ */d*_*S*_ *>* 1 only on foreground branches. The second alternative is closer to the truth, and therefore is chosen in a large number of cases, resulting in many false positives, and a low AUC. On the other hand, profile methods seem to suffer less from PPS, the effect of which can be compensated by increasing the effective population size *N*_*e*_. Effective population size *N*_*e*_ acts a scaling factor for the intensity of selection: as a result, observed amino acid frequencies are more representative of the actual fitness profile with higher values of *N*_*e*_, and constitute a stronger signal for profile methods.

### 4.6 Interpreting screens for changes in directional selection

The methods we discussed in this paper can be used to detect sites in alignments whose selection regime has changed coincidentally to a punctual transition event. Such transitions are typically changes in the environment, for example when a virus switches between hosts, and might also induce cases of convergent evolution (e.g the multiple transitions of mammals to the marine environment [Chikina et al., 2016]). In this context, these models can be used to give insights on the relation between the genotype and a given binary phenotype (e.g ancestral vs convergent, marine vs terrestrial, …). The fact that all methods except Multinomial are conservative, *i*.*e*., have low rates of false positives, indicates that the positives they output are likely to be worthy of further study.

The *d*_*N*_ */d*_*S*_ and profile methods that we discuss in this manuscript all make similar assumptions. Firstly, they can only handle a single phenotype or environmental condition at a time. This implicitly assumes that other phenotypes or conditions are unimportant for the evolution of the site under consideration. Such a strong assumption is likely to be incorrect in many cases: for instance a site may be important for several phenotypes, or its evolution may be more strongly associated to another phenotype or condition that has not been tested. Secondly, they assume that the evolution of the phenotype is known without uncertainty. *d*_*N*_ */d*_*S*_ approaches that can handle uncertainty in the evolution of the phenotype when reconstructing the evolution of gene sequences have recently been proposed, but remain to be extended to the site level [Halabi et al., 2021]. Thirdly, they rely on the comparison of two scenarios, one of which assumes homogeneity of the process across the phylogeny. In the *d*_*N*_ */d*_*S*_ method we consider, this means that the same *d*_*N*_ */d*_*S*_ parameter applies to the site throughout the phylogeny. In the profile methods we consider, this means that the same profile applies to the site throughout the phylogeny. This is likely to be incorrect: the site may be evolving inhomogeneously because of non-adaptive processes (e.g., CpG hypermutability or gBGC), or because it is correlated to unaccounted-for phenotypes or conditions. The use of homogeneous null scenarios can result in model confounding whereby an incorrect model is chosen in the absence of the true generating model [Jones et al., 2019]. This is what occurred in the gBGC simulations where the gBGC model generated data that was better fitted under our *H*_*a*_ model than under our homogeneous *H*_0_ model. However, our simulations of persistent positive selection show that profile methods are robust to this particular confounding process.

Our results show that a site found as positive with a profile method could result from a change in the direction (fig. 4) or intensity (fig. 5) of selection, as well as from a change in gBGC or PPS (fig. 6). At this stage, distinguishing between these processes requires looking at the profiles estimated by the method at the site. These profiles have been shown to be inferred accurately by several mutsel models [Spielman and Wilke, 2016]. Since codon-based methods do not perform better than amino acid-based methods in our hands, we suspect that the latter should also infer accurate profiles, although this will have to be verified in a future study. Given accurate profiles, one could distinguish between the different processes. Relaxation (respectively intensification) of selection should result in a flatter (resp. more heterogeneous) profile (fig. 1), which could be detected by computing its entropy and comparing it to the entropy of the other profiles at the site. gBGC should result in a shift towards GC-rich amino acids. PPS should result in a high amino acid diversity (large number of amino acids with non-zero frequencies).

### 4.7 Looking forward

The profile methods presented here have all been evaluated in the same setting, where the evolution of a site depends on two conditions that have been assigned to branches of a phylogeny. Not all phenotypes or conditions of interest can be known without uncertainty along a phylogeny, or can be accurately described by such a binary classification. Pelican can handle more than two conditions, but does not handle continuous annotations along a phylogeny, or uncertainty in the extant or ancestral states. Such extensions would be very useful. Similarly, the results show that accounting for gBGC in profile methods could be important. This could be done in codon models by following the approach that [Guéguen and Duret, 2018] used in *d*_*N*_ */d*_*S*_ models.

[Tamuri et al., 2014] and [Spielman and Wilke, 2016] showed that a penalized version of mutsel models performed better than the unpenalized version. We suspect that Pelican might also perform better with similar penalties. However, the use of penalized likelihoods would prevent us from relying on likelihood ratio tests to compute pvalues and detect positive sites. Instead, [Tamuri and dos Reis, 2021] relied on simulations to compute p-values, which is more ressource intensive and would compromise Pelican’s scalability. More work is needed to investigate the benefits of using penalization in Pelican, and, if any, come up with a fast method to compute p-values or scores. Such a method might also improve on the LRT that we have used here, as we saw that tinkering with its degrees of freedom improved the performance of the method in some cases (sup. fig. S13).

Overall, the results show that profile methods constitute a solid alternative to *d*_*N*_ */d*_*S*_ methods to screen for substitutions associated to changes in a phenotype or condition of interest. This opens new possibilities to better understand the link between a substitution, the structure of the protein where it occurs, and the phenotype or condition to which it is correlated. The amino acid profiles inferred by a profile method at a site can be used to investigate the effect that having a high fitness or a low fitness amino acid has on a protein structure, in a particular condition. Profile methods could thus pair very well with the recent improvements in protein structure prediction [Jumper et al., 2021] to yield new insights into the molecular basis of adaptation.

## 5 Conclusion

In this paper we evaluated on simulations a series of methods aiming to detect changes in selective pressures in coding sequences along a phylogeny. We found that some profile methods compare favourably to a commonly used *d*_*N*_ */d*_*S*_ method, both in terms of power and in terms of speed, including in the presence of confounding factors. In particular, profile methods can readily distinguish changes in directional selection from persistent positive selection, something that the *d*_*N*_ */d*_*S*_ method we tested cannot do. Among profile methods, we found that Pelican, a method operating at the amino acid level, can be used to detect selective pressure changes efficiently. This makes genome-wide searches for sites correlating with a phenotype or condition of interest doable on a single computer within a few days.

Further extensions of Pelican are envisioned, for example to handle continuous phenotypes. Integrating the effect of gBGC in the model would also be a major improvement, as we have found that it has a strong confounding effect on the detection of selection.

## 6 Methods

### 6.1 Detection of *ω* variations using codeml

We used the `codeml` tool from the PAML package to detect variations of *d*_*N*_ */d*_*S*_ as a proxy for variations of selective pressure, as was done in [Thiltgen et al., 2017]. Branch lengths were re-estimated by `codeml`. `codeml` was configured to use the branch-site model A [Zhang, 2005, Yang, 2005]. This model assumes there are three categories of sites in the alignment, whose proportions are estimated. Categories `0` and `1` have a homogeneous *ω* value throughout the phylogeny. Category `2` has one *ω* value estimated per branch condition: on background branches, the *ω* is between 0 and 1 (subcategory `2a`), characteristic of purifying selection, or at 1 (sub-category `2b`), characteristic of neutral evolution. On foreground branches, *ω ≥* 1, characteristic of neutral or positive selection. A site is declared “positive” if it belongs to this category `2`. The probability for each site to be positive as inferred by the method was computed from the Bayes empirical Bayes probabilities, resulting from running `codeml` with parameter `fix omega = 0` and summing up the probabilities to belong to categories `2a` and `2b` in the model.

### 6.2 Multinomial method

The multinomial method models each site of an alignment as a collection of independent categorical variables, thus completely ignoring the phylogeny. It compares two models using a likelihood ratio test (LRT), the first one assumes a single probability vector of length 20 (one frequency per each amino acid), the second a pair of vectors, one for each condition. Computing a *p*-value is however difficult in our setting, as at a given site, most of the amino acids are not observed and as a consequence their frequency estimated by maximum likelihood is zero, and thus lies at the boundary of the parameter space. In that case the usual convergence of the likelihood log-ratio to a *χ*^2^ distribution known as Wilks theorem does not hold. While there exists literature on the subject (see [Mitchell et al., 2019] for a recent result), existing results are difficult to apply. We reused a heuristic we found in [Tamuri et al., 2009], consisting in approximating the likelihood log-ratio distribution under the null by a *χ*^2^ distribution with number of degrees of freedom equal to the number of amino acids observed at the leaves of the tree minus one.

### 6.3 Pelican : improvements on TDG09

Pelican is a reimplementation of the TDG09 method, originally published by [Tamuri et al., 2009]. TDG09 relies on a site-independent model of amino acid sequence evolution and the WAG exchangeability matrix. The model involves two kinds of parameters : stationary distributions of amino acids and branch scale.

Inference of selective pressure shifts is based on the postulate that stationary distributions of amino acids reflect the fitness profile in a condition (e.g. foreground or background). In a similar way to the multinomial method, the likelihoods of two models are compared using the LRT procedure, where one model assumes a single stationary distribution of amino acids shared between both conditions, and the other model assumes a specific stationary distribution per condition.

Parameters of the model, such as stationary distributions and branch scale, are optimized to maximum likelihood using the Nelder-Mead algorithm [Nelder and Mead, 1965]. We implemented an alternative approach using automatic differentiation, made available through the PyTorch library [Paszke et al., 2019], that converges to the same optima as the Nelder-Mead implementation. This alternative optimisation algorithm is currently not used, but might be useful in future extensions of the method.

Pelican is implemented in the OCaml language [Leroy et al., 2021]. The underlying mutationselection model implementation takes advantage of LAPACK [Anderson et al., 1999] bindings for fast linear algebra computation, and optimisations for transition matrices exponentiation through diagonalisation [Yang, 2006]. Pelican is available at https://gitlab.in2p3.fr/phoogle/pelican.

### 6.4 Simulations

In all our experiments, simulations were used to generate amino-acid or codon alignments with a constant number of sites *N* = 10 000. The simulator was configured to generate 90% of *H*_0_ sites (no changes in selective pressure) and 10% of *H*_*A*_ sites (different selective pressure between background and foreground condition). Simulations were done using a general time-reversible (GTR) mutation-selection model at the codon level. The model allows for two different regimes : one modeling selection in the background condition, and the other in the foreground condition. Selective pressure changes on *H*_*A*_ sites are simulated using either the foreground or background regime, depending on the condition of each branch in the phylogenetic tree. *H*_0_ sites are generated using only the background regime, indicating no change in the selective pressure through the tree for these sites. Each regime is represented as a matrix of substitution rates between codons, which can be run along the phylogeny using Gillespie’s algorithm [Gillespie, 1976].

The substitution rates are the result of a mutation probability and a relative fixation probability, which depends on a selection coefficient associated with the transition to the mutated state. Mutation probabilities for the GTR model of nucleotide substitutions are based on exchangeabilities drawn from a *Gamma*(1, 1) distribution, and equilibrium frequencies from a *Dirichlet*(10, 10, 10, 10) distribution, and are shared across sites. The selection coefficient *S* (eq. 1) is defined as the difference in fitness between the ancestral state *X* and the mutated state *Y* in a condition *c*.

The relative fixation probability *u*(*S*) for a mutation is computed from the selection coefficient *S* as per [Kimura, 1983]:

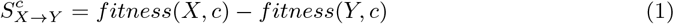

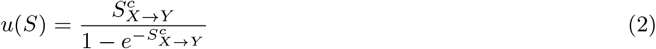

Fitness values are determined from amino acid frequency profiles, which are randomly picked at each site from a set of 263 preset profiles [Rey et al., 2019] for each condition. These frequency profiles are transformed into fitness profiles by multiplying them by a factor *Ne* = 4. As a result, values 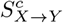 are between *−*4 and 4.

Codon substitution rates *σ* are the product of mutation rates *µ* and the relative probability of fixation :

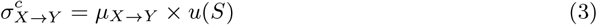

### 6.5 gBGC simulation

GC-biased gene conversion (gBGC) acts as a fixed increase in fixation probability for mutations from either A or C nucleotides to G or C; conversely it is modeled as a probability decrease when mutating the other way around. We included GC-biased conversion in our simulation model as a bias term in the selection coefficient *S*:

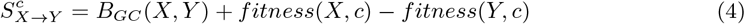

Based on [Glémin et al., 2015], we chose an intensity of *B*_*GC*_ = 10, that is applied on fore-ground branches, which is a strong effect for this process. Transition rates were not affected on background branches. This way, in *H*_*A*_ sites, the change in selective pressure between background and foreground branches that has to be detected is driven both by the shifted fitness profile, and the effect of gBGC. In *H*_0_ sites, gBGC affects foreground branches.

### 6.6 CpG simulation

CpG hypermutability is introduced in the simulation model as a scaling factor *ρ* for the mutation probability:

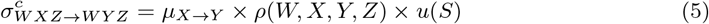

where *W* and *Z* are the states at the surrounding sites. This context is necessary because CpG dinucleotides can occur across two codons. As a consequence, the evolution of a whole sequence is not site-independent anymore, which led us to develop a dedicated Gillespie simulator. CpG hypermutability only occurs on methylated CpG dinucleotides, and induces an increased probability of mutation from C to T in this context (or G to A on the reverse strand). We assume that any CpG dinucleotide in our simulation is methylated. If the conditions for hypermutability are not verified when comparing changes from *X* to *Y*, or the current branch is background, *ρ*(*W, X, Y, Z*) = 1 and has no effect. Otherwise, on foreground branches, we set *ρ*(*W, X, Y, Z*) = 10 based on [Meunier et al., 2005], both on *H*_*A*_ and *H*_0_ sites.

### 6.7 Simulation of persistent positive selection

PPS is introduced in the simulation model as a constant increasing the fitness of all other amino acids except the current one [Tamuri and dos Reis, 2021]. This is achieved by modifying equation 1 as:

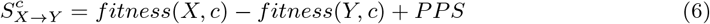

where *PPS ≥* 0 is a constant and describes the strength of positive selection. To simulate data, we relied on two parameter settings. In the first setting, we simulate sequences under a mild selection strength, setting *Ne* = 4 and *PPS* = 2. This setting ensures that differences in amino acid fitnesses are between *−*4 and 4, as in the rest of the manuscript. In the second setting, we simulate under a strong selection regime, with *Ne* = 10 (*i*.*e*., differences in amino acid fitnesses between *−*10 and 10), and *PPS* = 10. This second setting resembles parameter values observed on the sites showing the strongest positive selection in [Tamuri and dos Reis, 2021], and is also similar to their own simulation settings. *H*_*A*_ sites were simulated with different profiles for background and foreground branches, and *H*_0_ sites were simulated with PPS running both on background and foreground branches.

## 7 Code and data availability

Source code to reproduce the results in this paper is publicly available at

https://gitlab.in2p3.fr/phoogle/spcd-benchmark.

The implementation of Pelican is also made available at

https://gitlab.in2p3.fr/phoogle/pelican.

Plots were produced in R [R Core Team, 2021] using the packages *ggplot2* [Wickham, 2016] and

*ggtree* [Yu, 2020].

## 8 Acknowledgements

We thank Nicolas Lartillot and Anamaria Necşulea for their comments on an early version of the manuscript and fruitful discussions. The research presented here was funded in part by the Convergenomix project (ANR-15-CE32-0005). L.D. was supported by a PhD fellowship from the Université Lyon 1 Claude Bernard.

