## Supplementary material for "Evaluation of methods to detect shifts in directional selection at the genome scale"

March 2022

#### S1 Synthetic trees

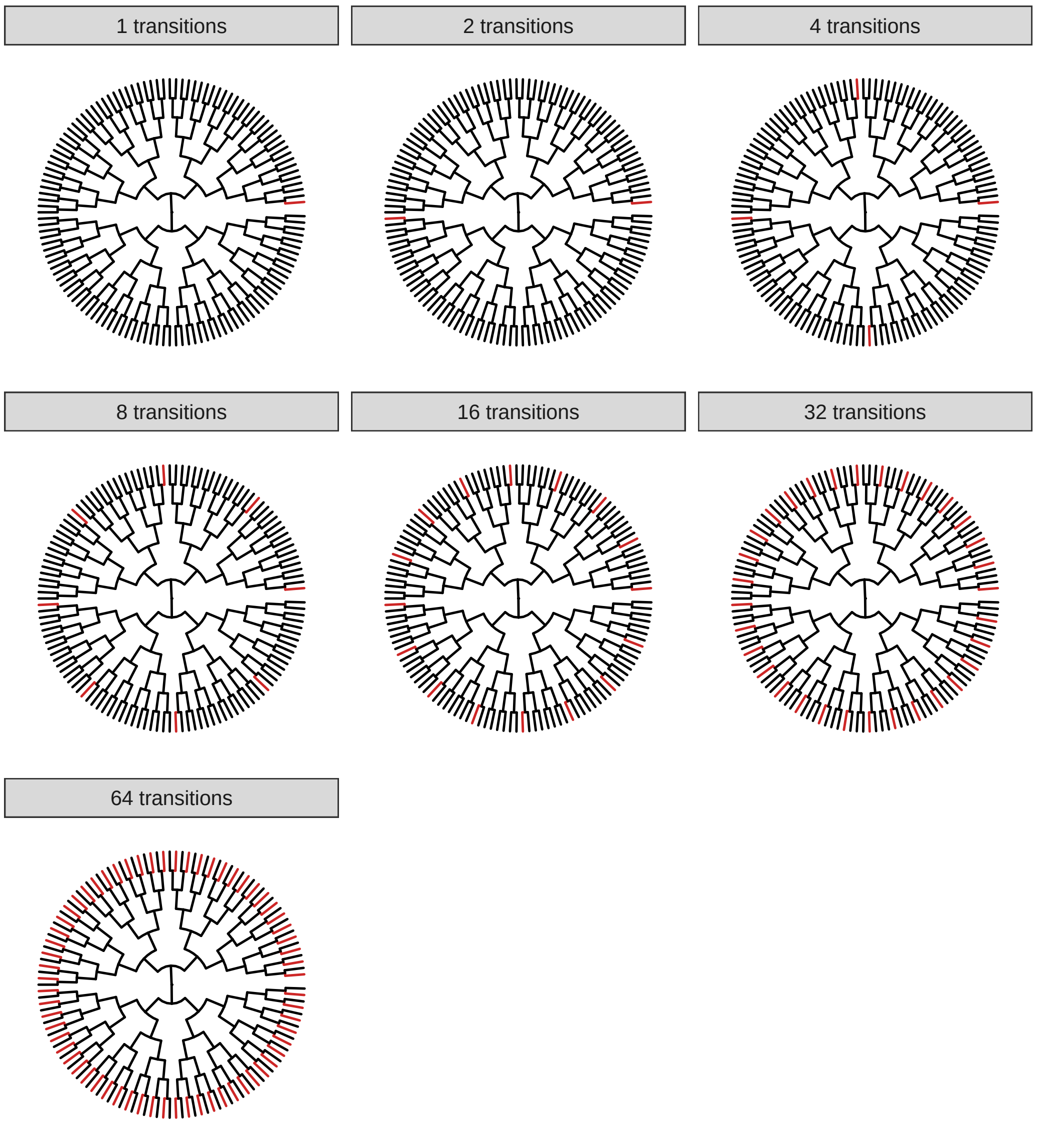


Figure S1: Synthetic trees with variable number of transitions on terminal branches

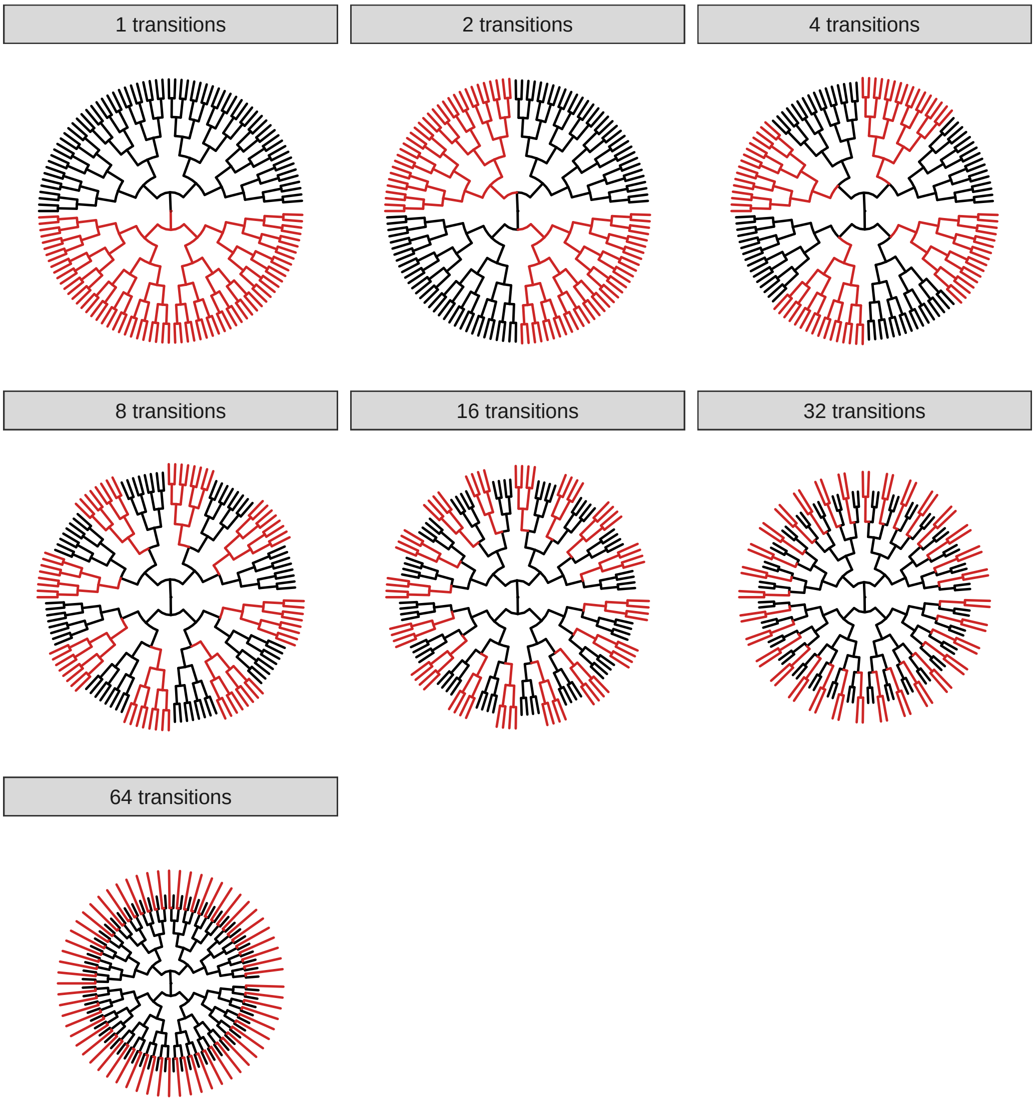


Figure S2: Synthetic trees ensuring a constant amount of time spent in a condition

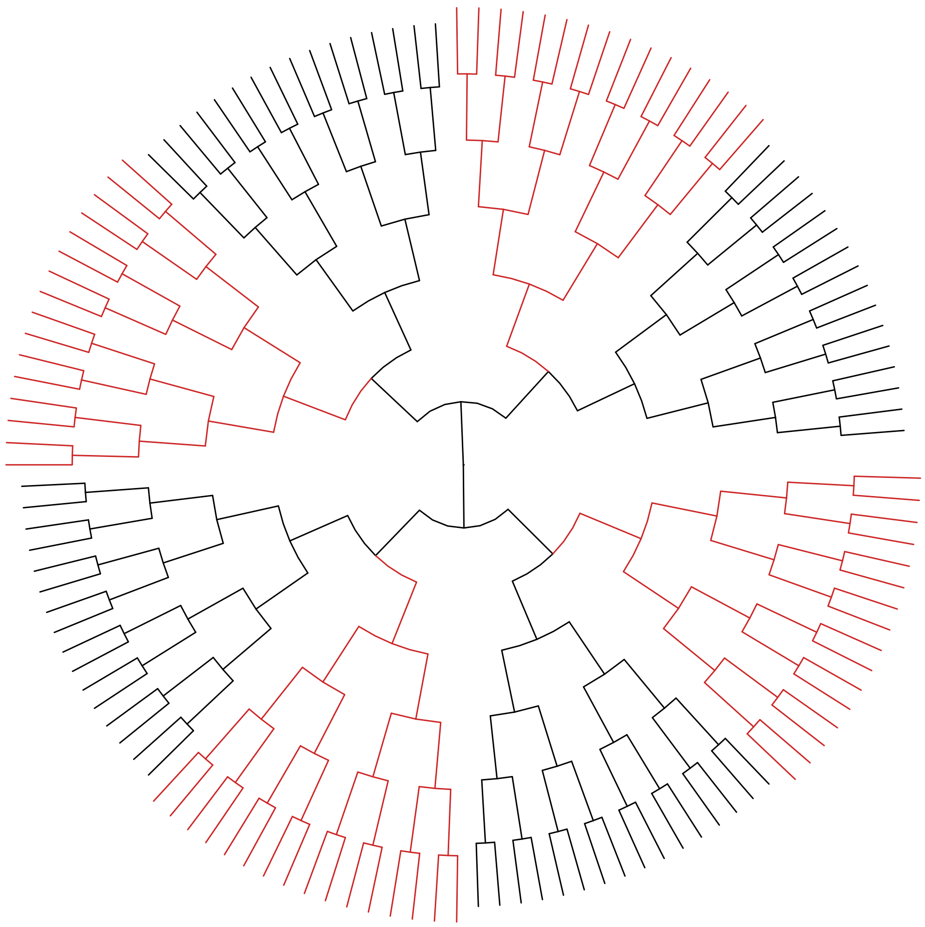

Figure S3: Tree topology and annotation used in the experiment where variation in branch lengths is investigated.

#### S2 Empirical phylogenies

Tree scale is given in expected number of codon substitutions.

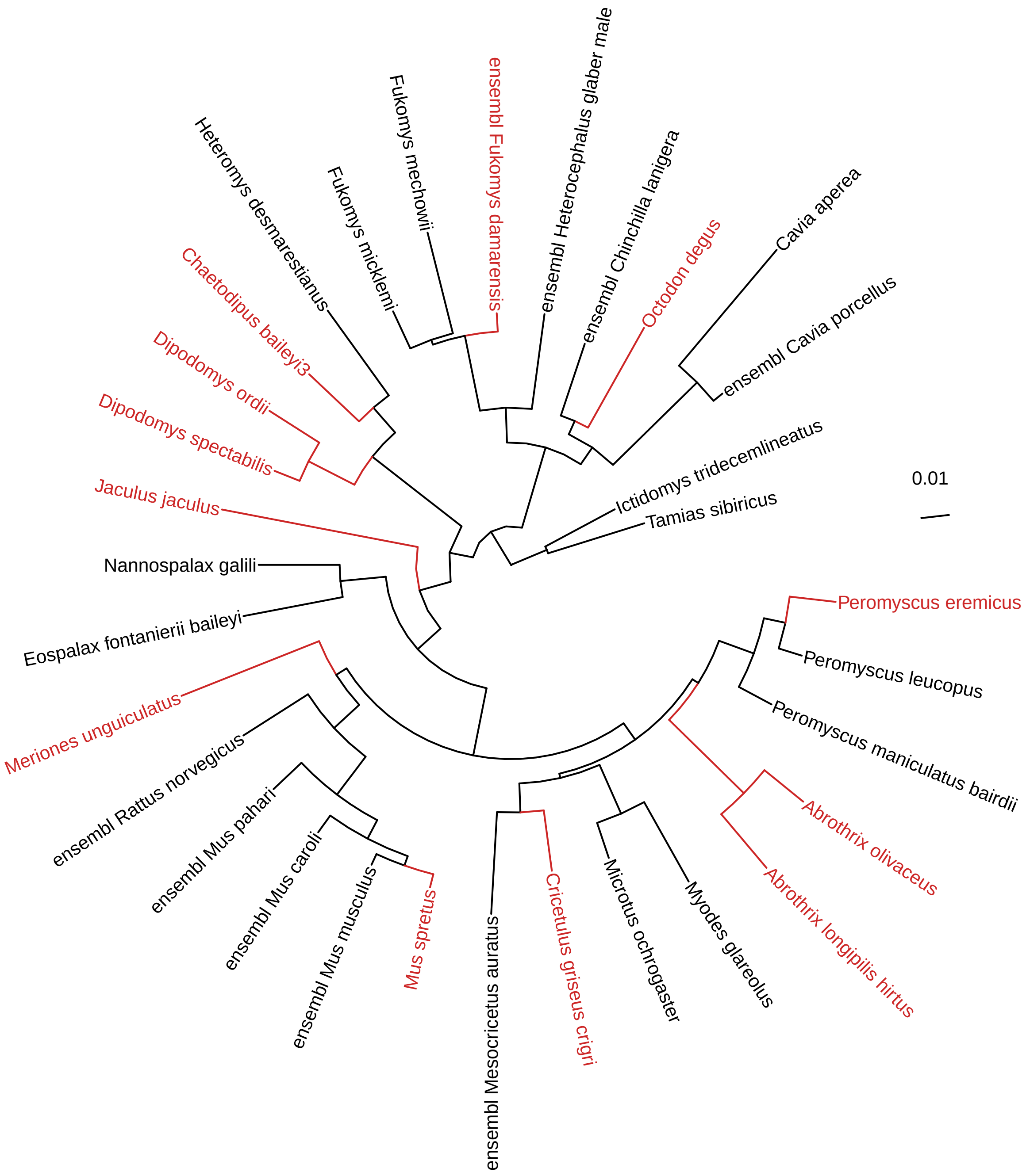


Figure S4: Rodents tree [Rey et al., 2019]

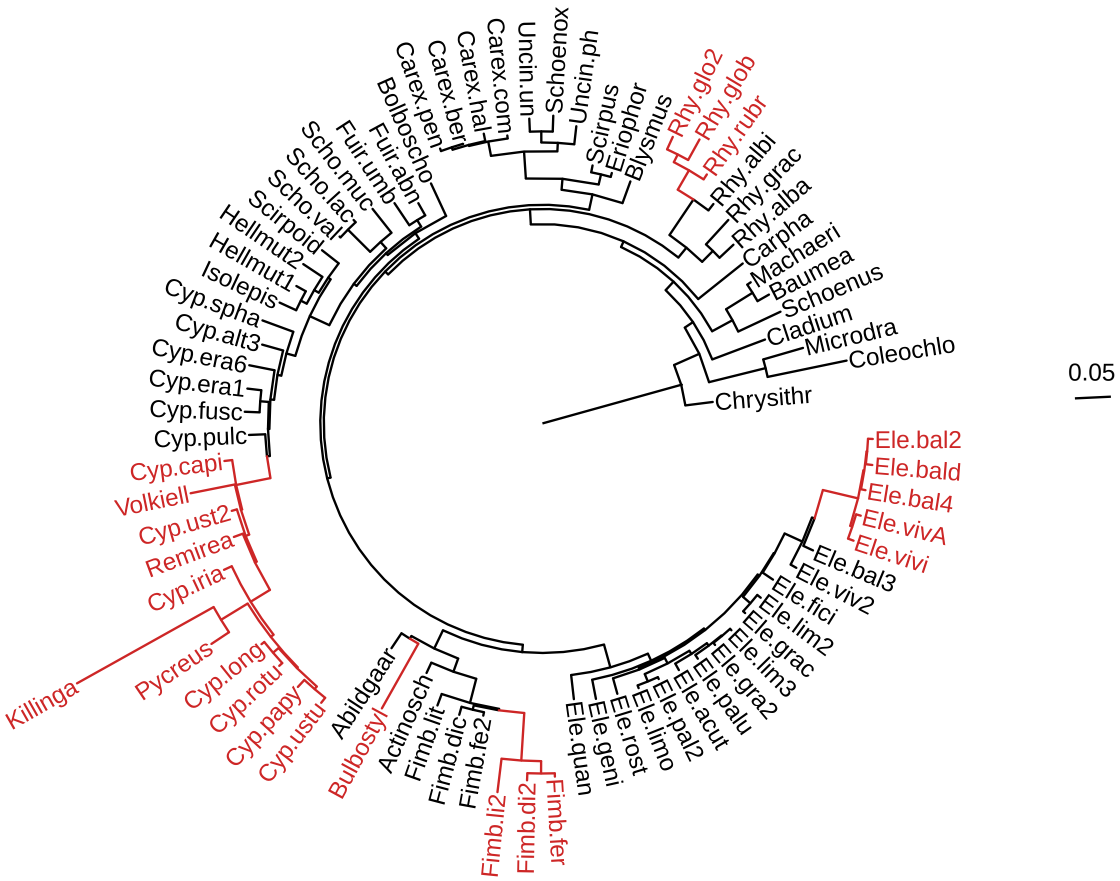


Figure S5: *Cyperaceae* tree [Besnard et al., 2009]

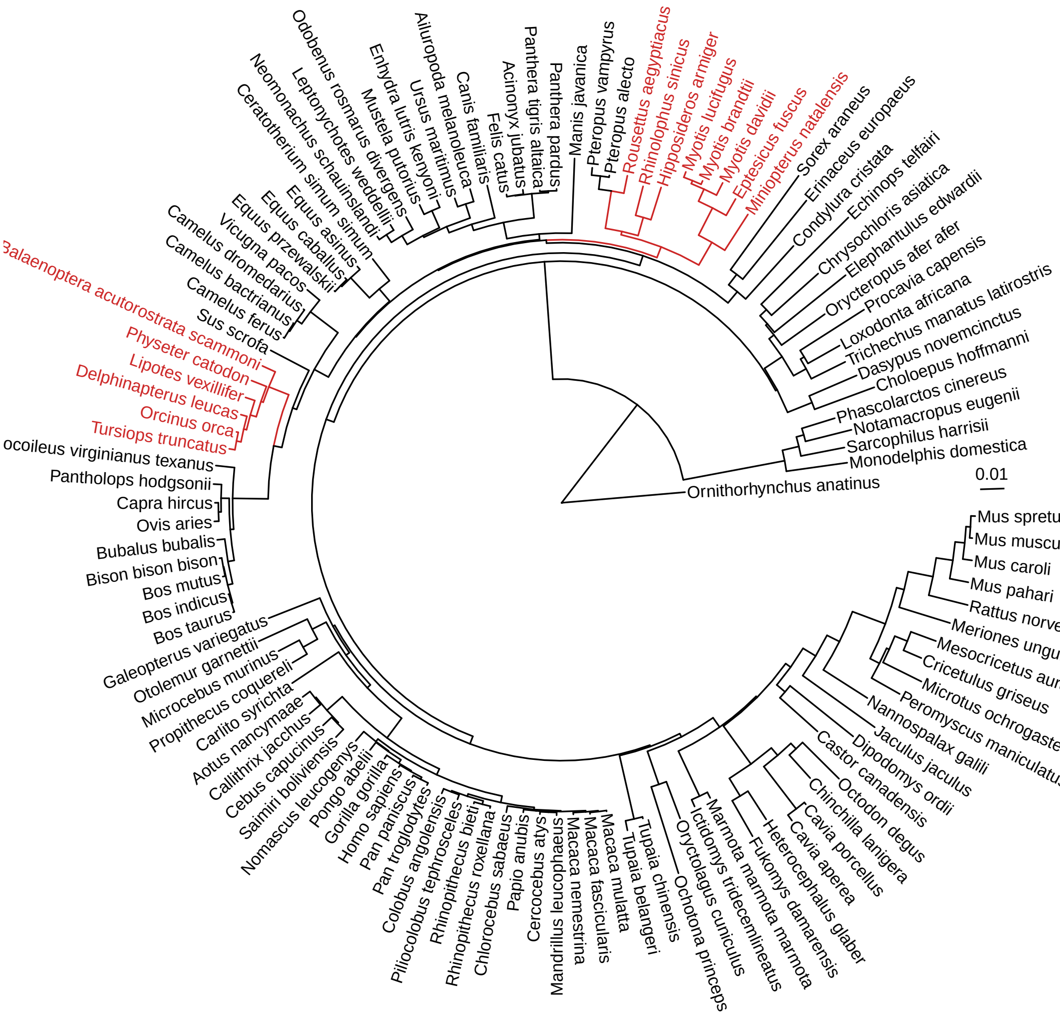


Figure S6: Echolocation tree [Scornavacca et al., 2019]

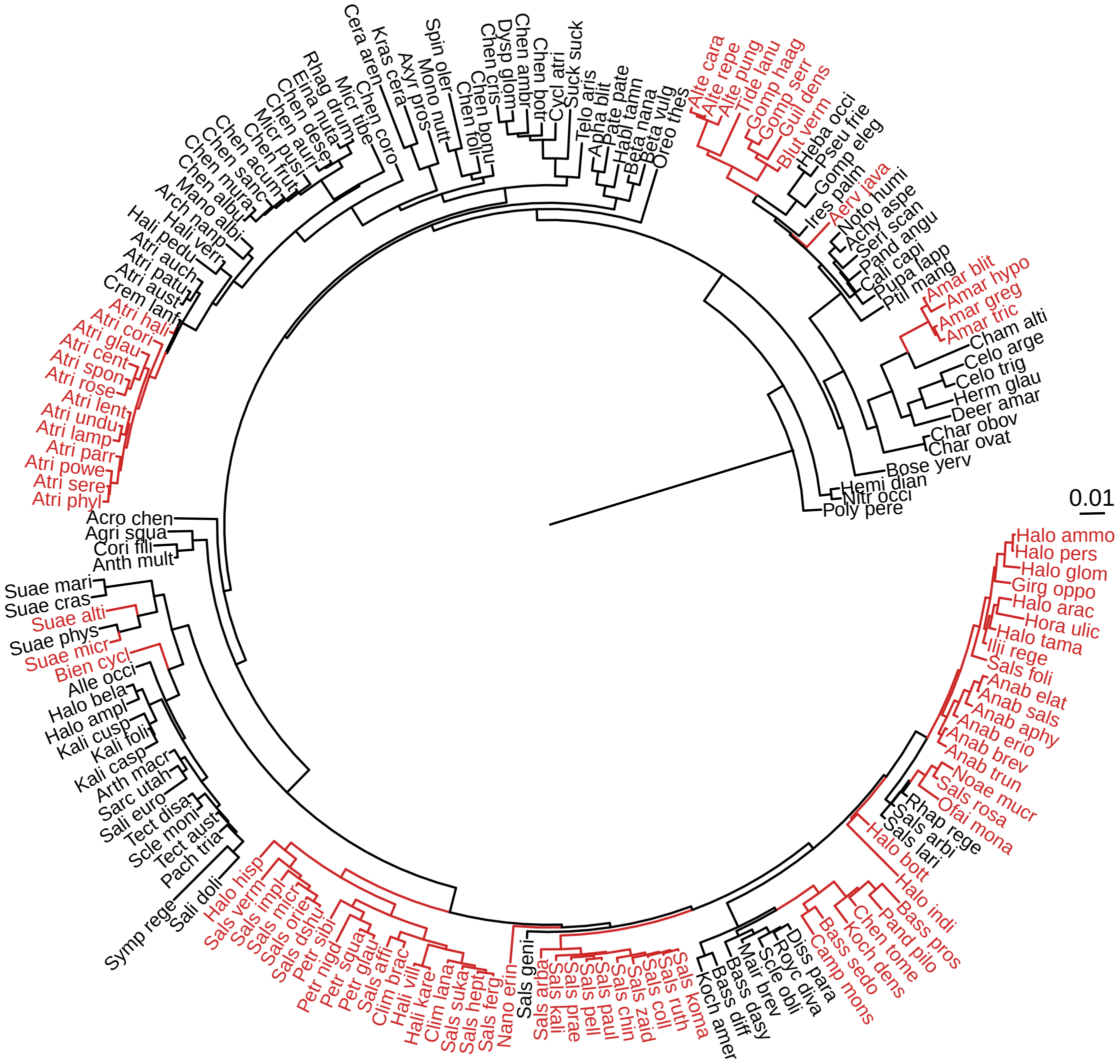


Figure S7: *Amaranthaceae* tree [Kapralov et al., 2012]

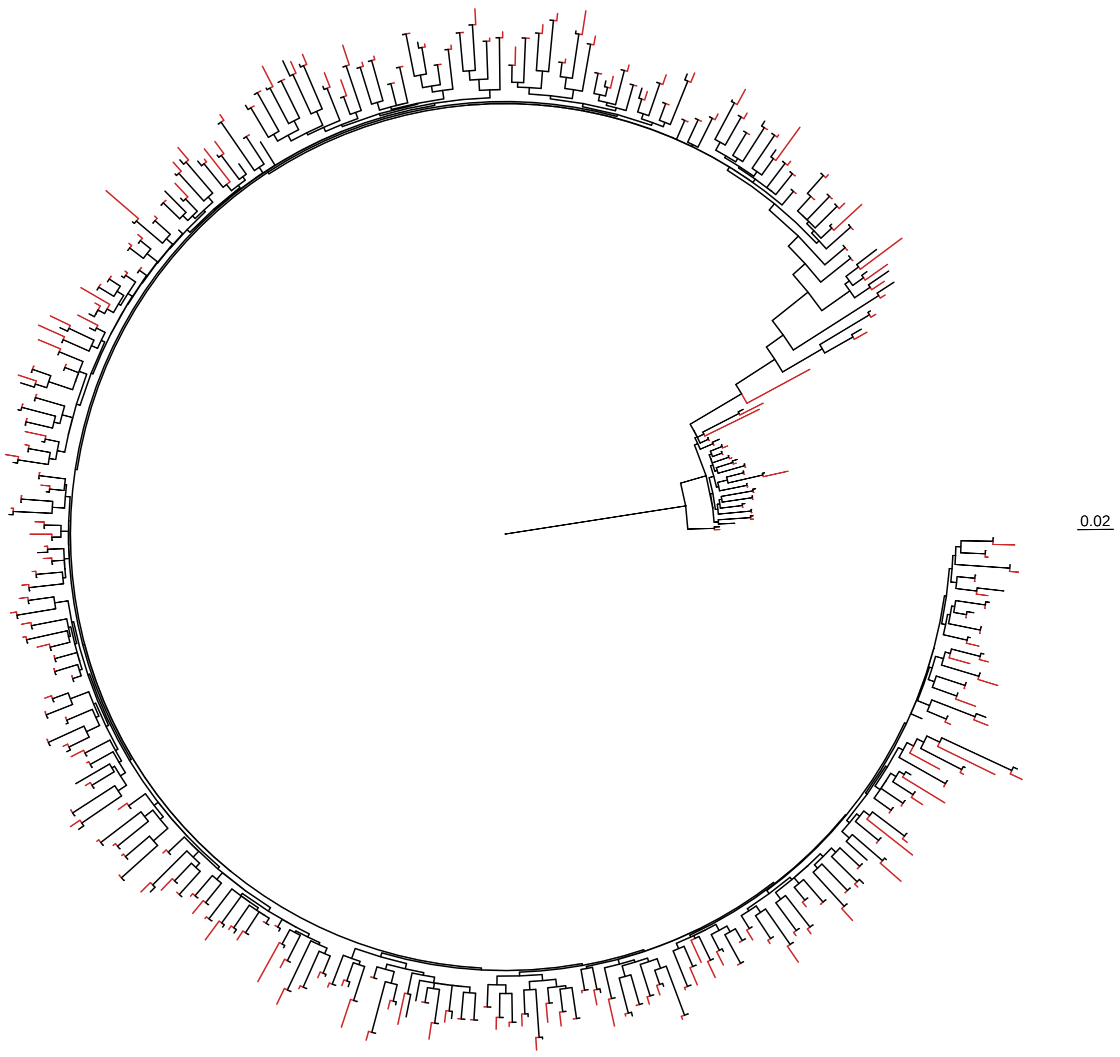


Figure S8: HIV Reverse-transcriptase tree [Murrell et al., 2012]

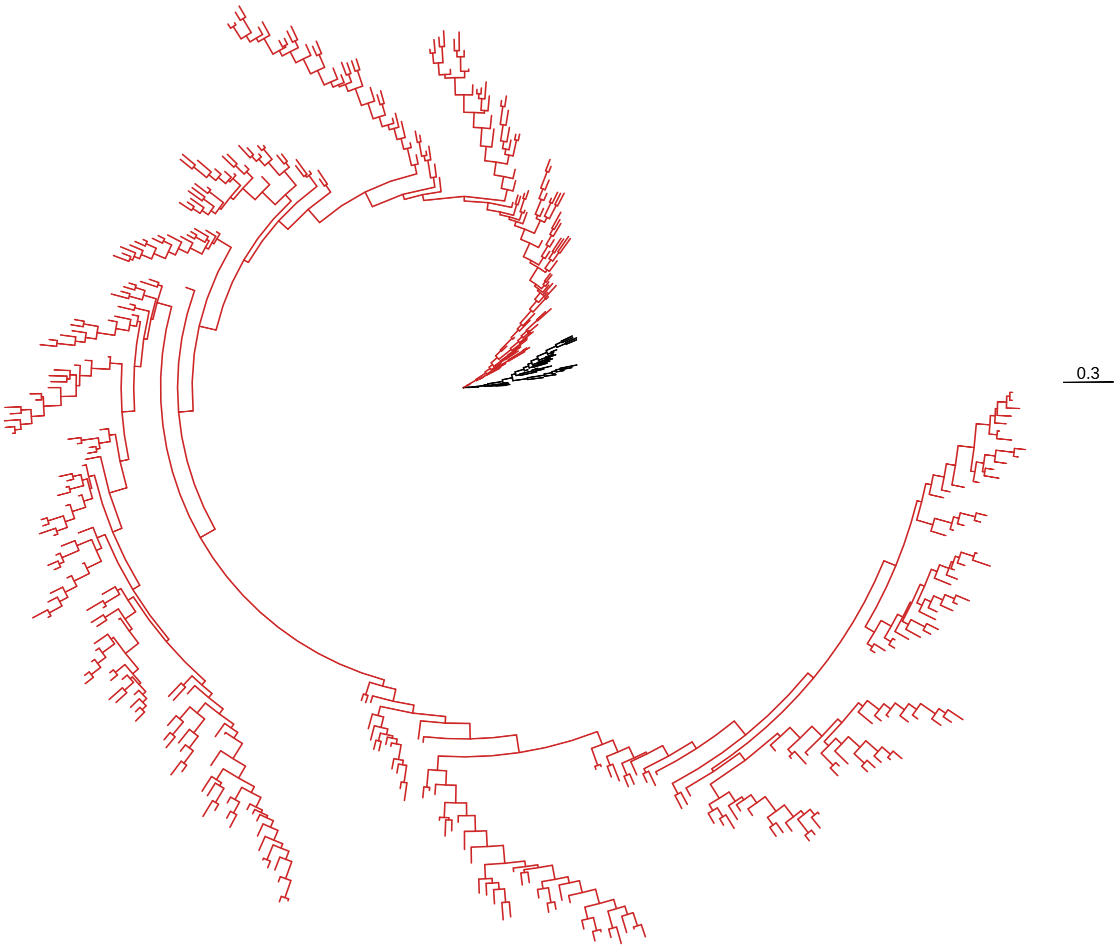


Figure S9: Influenza H1 segment [Tamuri et al., 2009]

#### S3 Study of the calibration of the methods

| Dataset | Diffsel | Gemma | Multinomial | PCOC | Pelican | TDG09 | codeml |
|---|---|---|---|---|---|---|---|
| *Cyperaceae* | 0.00 | 0.02 | 0.07 | 0.00 | 0.02 | 0.03 | 0.00 |
| *Amaranthaceae* | 0.00 | 0.00 | 0.05 | 0.00 | 0.01 | 0.02 | 0.00 |
| Rodents | 0.00 | 0.00 | 0.00 | 0.00 | 0.00 | 0.01 | 0.00 |
| Echolocation | 0.00 | 0.01 | 0.02 | 0.00 | 0.01 | 0.02 | 0.00 |
| HIV | NA | 0.00 | 0.00 | NA | 0.01 | 0.01 | 0.00 |
| Influenza | NA | 0.04 | 0.23 | 0.01 | 0.05 | NA | 0.02 |

Table S1: Observed false positive rate at the 0.05 threshold. A well calibrated method should yield a proportion of 0.05 false positives.

| Dataset | Diffsel | Gemma | Multinomial | PCOC | Pelican | TDG09 | codeml |
|---|---|---|---|---|---|---|---|
| *Cyperaceae* | 0.00 | 0.08 | 0.24 | 0.00 | 0.06 | 0.12 | 0.00 |
| *Amaranthaceae* | 0.00 | 0.02 | 0.24 | NA | 0.02 | 0.10 | 0.00 |
| Rodents | 0.00 | 0.02 | 0.01 | 0.00 | 0.02 | 0.11 | 0.00 |
| Echolocation | 0.00 | 0.05 | 0.10 | 0.00 | 0.05 | 0.10 | 0.00 |
| HIV | NA | 0.00 | 0.00 | NA | 0.02 | 0.04 | 0.00 |
| Influenza | NA | 0.07 | 0.41 | 0.01 | 0.09 | NA | 0.25 |

Table S2: Observed false positive rate at the 0.05 threshold, after removing constant sites from the data. A well calibrated method should yield a proportion of 0.05 false positives.

#### S4 Benchmark results with confounding factors

Evaluations on empirical phylogenies in the presence of confounding factors are presented here. They include persistent positive selection (PPS) and non selective forces : GC-biased gene conversion (gBGC) and CpG hypermutability. HIV and Influenza datasets were excluded from this analysis, because of their larger size and the associated computational costs.

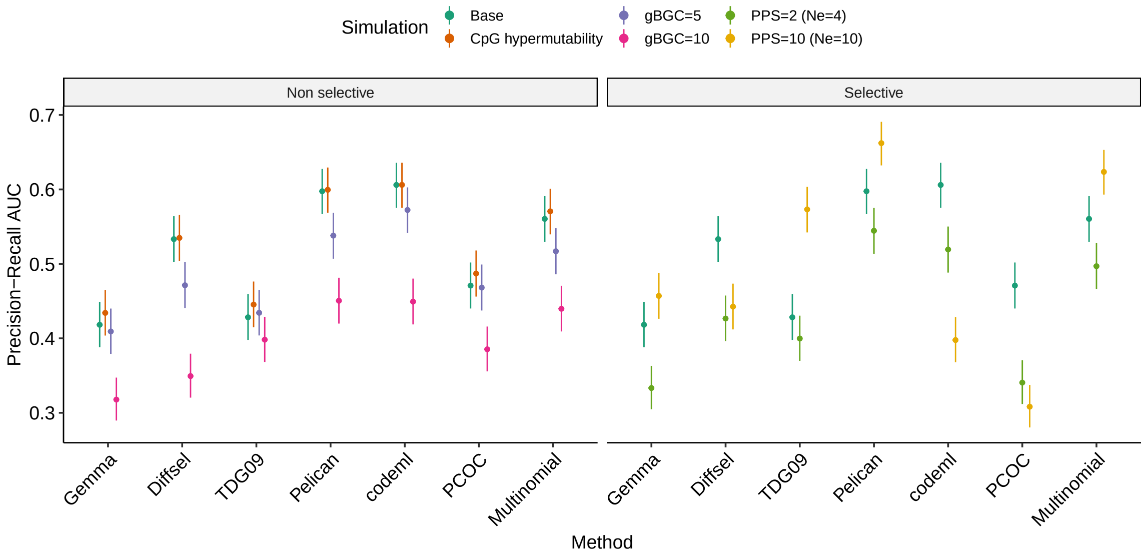


Figure S10: Precision-recall AUC for all methods in the presence of confounding factors measured on the Rodents dataset.

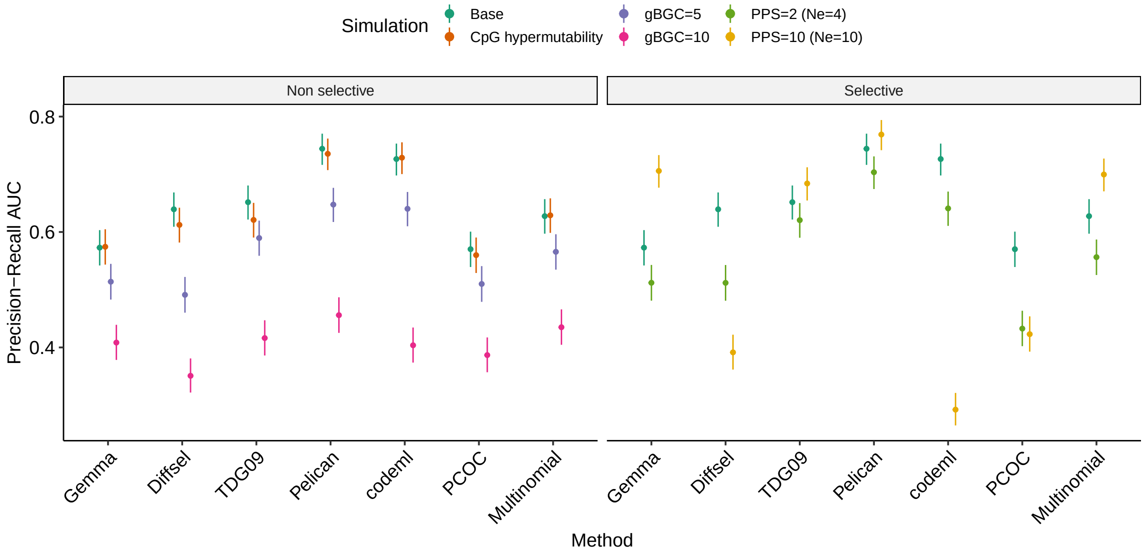


Figure S11: Precision-recall AUC for all methods in the presence of confounding factors measured on the *Cyperaceae* dataset.

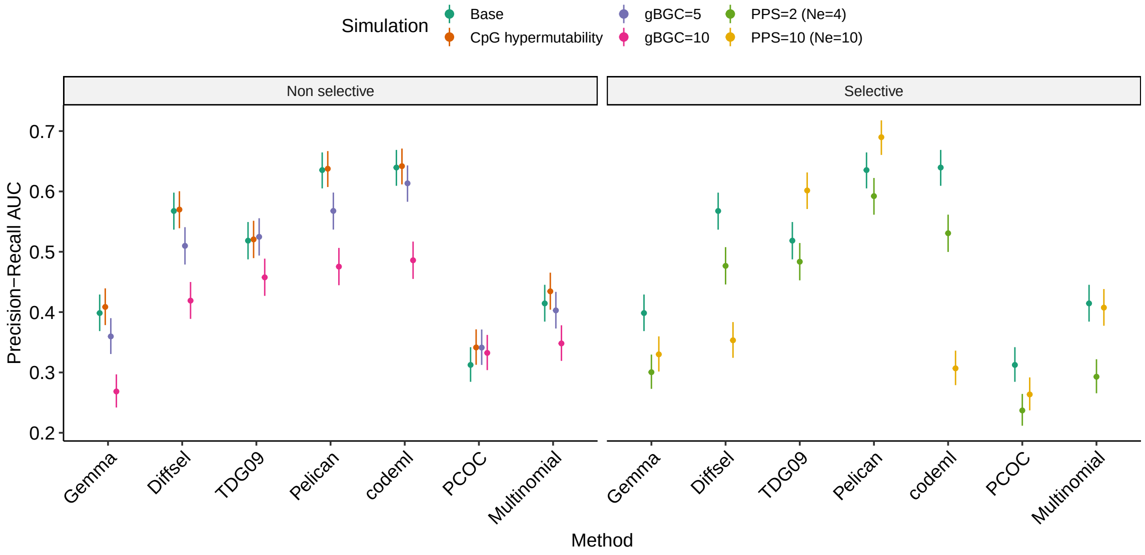


Figure S12: Precision-recall AUC for all methods in the presence of confounding factors measured on the *Amaranthaceae* dataset.

### S5 Evaluation of Pelican using different degrees of freedom in the LRT

In TDG09 [Tamuri et al., 2009], the degrees of freedom for the LRT were computed as the number of amino acid types observed at one site minus one, corresponding to the number of additional adjustable parameters in the alternative ($H_A$) model compared to $H_0$. Our implementation of this model, named Pelican, uses the same specification for the computation of degrees of freedom.

In response to the observation that Pelican shows strongly decreased performance for detecting relaxations of selective pressure (main fig. 5), we investigated a different specification for the computation of the degrees of freedom as the difference between the sum of the numbers of amino acid types observed in each condition, and the total number of amino acid types observed at one site:

$$\text{df} = \left( \sum_c |AA_c(site)| \right) - |AA(site)| \qquad \text{where } c \text{ is a condition}$$

We name this alternative specification `Pelican alt`. Both specifications were evaluated on all empirical phylogenies (sup. fig. S13).

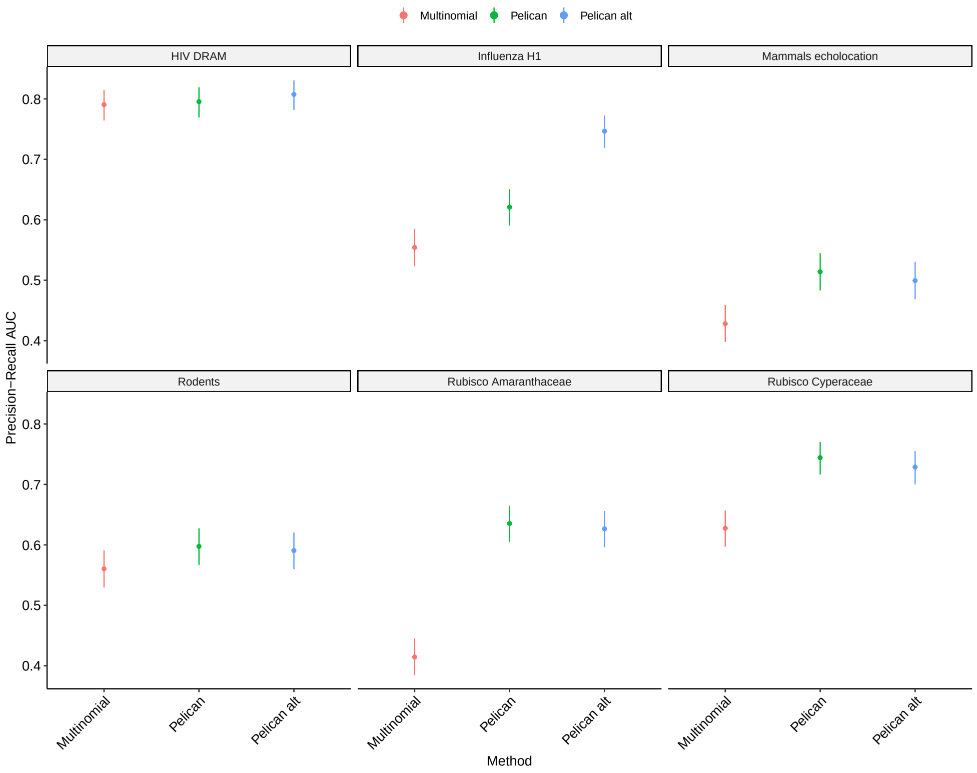


(a) Differential selection

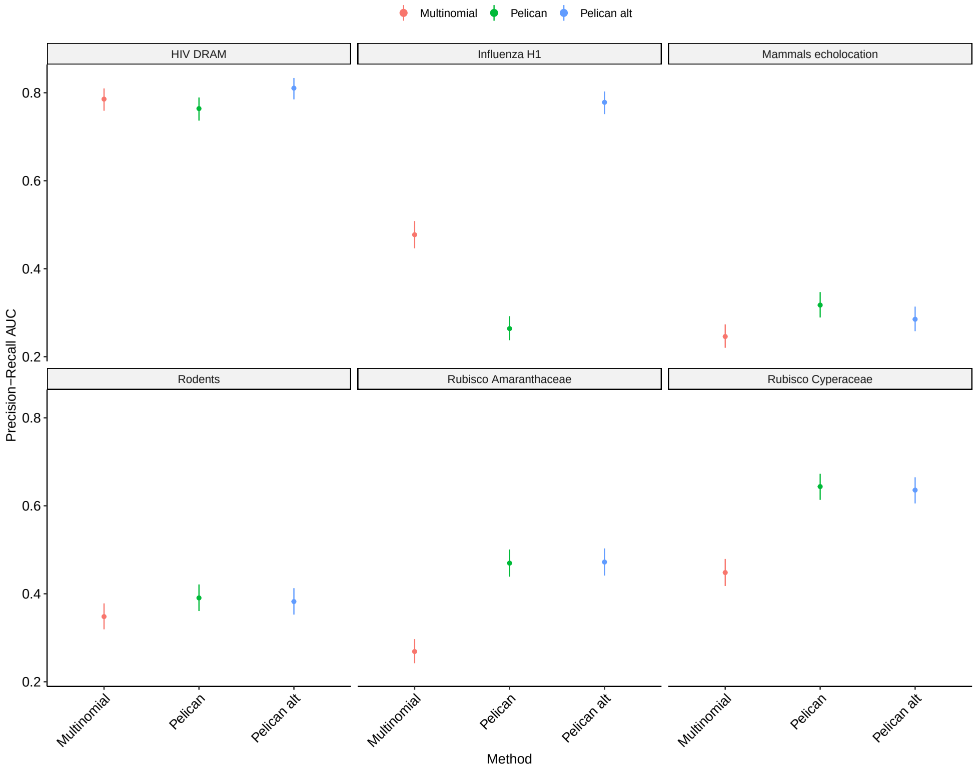


(b) Relaxed selection

Figure S13: Evaluation of Pelican using two different specifications for the computation of degrees of freedom in the likelihood ratio test. `Pelican` is the version that was evaluated throughout the main manuscript, and uses the original specification from [Tamuri et al., 2009]. `Pelican alt` uses a different specification that is defined in the manuscript (Material and methods).

The alternative specification appears to be a good trade-off, as performance is slightly decreased on some datasets (e.g Echolocation), but strongly increased on others (e.g Influenza, HIV). However, it differs from the specification in the original implementation of the model [Tamuri et al., 2009], and from the usual specification of a LRT [Wilks, 1938], and needs to be more thoroughly tested. Altogether, these results suggest that further improvements can be made to Pelican by improving how the LRT is computed.
